# Flexible Perception of Tactile Cues in Multiple Reference Frames

**DOI:** 10.1101/2023.11.10.566625

**Authors:** Himanshu Ahuja, Sabyasachi Shivkumar, Catalina Feistritzer, Ralf M. Haefner, Gregory C. DeAngelis, Manuel Gomez-Ramirez

## Abstract

Leading models of somatosensory processing posit that integration of tactile and proprioceptive cues is essential for touch perception. Yet, such integration is not universally beneficial. While posture-dependent remapping of tactile signals is critical for guiding hand actions, discrimination of certain tactile features is enhanced when touch is represented independently of hand posture. How the brain flexibly controls the integration of tactile and proprioceptive cues based on task demands remains unclear, as most studies have only examined touch perception within a single reference frame. Here, we studied how reference frame demands shape touch perception in humans making motion judgements on a finger, while varying hand posture. Participants were cued to report motion direction relative to the finger or the sternum (i.e., the body’s midline). We found that tactile and proprioceptive cues are integrated in a task-specific manner, with hand posture biasing judgments only in the Sternum-centric task. Further, we observe that reaction times are faster for Sternum-centric judgments, with accumulation-to-bound models indicating that this advantage is driven by lower decision thresholds. Finally, we developed a Bayesian computational framework that formalizes how tactile motion signals are transformed across hand postures, advancing our understanding of how the brain maps sensory information from skin-to body-centered coordinates. Together, these findings demonstrate that reference frame task demands shape both the transformation of tactile information and speed of perceptual decisions, within a Bayesian computational framework for task-dependent coordinate transformations.

## Introduction

Humans have exceptional hand dexterity that supports many haptic behaviors such as tool use, object sensing, social touch, and others. This ability commences with specialized mechanoreceptors and proprioceptors that transmit signals encoding different features of the object to the brain, where they combine to form a holistic representation of the object^1–5^. A key function of the somatosensory system is to relay tactile feedback to motor areas to guide adjustments for sensing and grasping, but these sensory signals should be mapped into coordinate frames that align with the sensorimotor demands of the behavior. For example, perceiving how an object moves relative to the body requires integrating cutaneous motion with proprioceptive signals and transforming this information into an egocentric (trunk/sternum-or head-centered) reference frame to maintain perceptual constancy across changes in hand posture. In contrast, adjusting the grasp when an object begins to slip requires representations in a hand-centered (or finger-centered) reference frame that encode the spatial relationships between the digits (e.g., the relative positions of the thumb and fingertips during a pinch). Similarly, to perceive non-spatial features such as texture or temperature, it is advantageous to represent tactile signals independently of hand posture (e.g., in skin-centered coordinates). Thus, rather than committing tactile signals to a single coordinate system, the brain appears to flexibly represent touch in the reference frame that is most useful for the task. However, the computational mechanisms underlying this flexibility remain poorly understood, in part because most studies have examined tactile and proprioceptive interactions using tasks that require judgments within a single reference frame^6–19^. In particular, it is unknown whether tactile and proprioceptive signals are combined the same way regardless of task (i.e., with fixed weights), or whether the brain selectively modulates their integration according to task demands (i.e., using task-adaptive weights). For instance, when a task does not require integration of tactile and proprioceptive inputs, does the brain optimally segregate the two modalities, or is it constrained in how they are combined? Conversely, when perception benefits from integration, does the brain do so optimally, remapping tactile inputs based on proprioception, or are there systematic biases in the integration?

A previous study showed that tactile motion perception is modulated by proprioception, with perceptual biases predicted by the relative distance between the finger and head^20^. However, participants were not cued to use a specific reference frame and likely defaulted to head-centered judgments based on the task instructions. Thus, it remains unknown whether the brain implements an optimal weighting strategy for tactile and proprioceptive signals, or whether it employs other schemes such as fixed weighting or intermediate weights that partially change with task demands. To address this gap in knowledge, we conducted a series of human psychophysical studies that explicitly cued motion discrimination in different reference frames (relative to the sternum or to a finger) while systematically varying hand posture, allowing direct comparisons of how proprioceptive and tactile signals are integrated across reference frame contexts. We found that humans flexibly represent tactile motion in a task-specific reference frame, with proprioception modulating perception only in the Sternum-centric task. We also found that judgements are faster in the Sternum-centric task, consistent with a perceptual readout mechanism that favors touch encoded in body-centered coordinates. Accumulation-to-bound models reveal that the faster reaction times (RTs) in the sternum-centric task are driven primarily by lower decision thresholds, indicating a difference in decision policy consistent with a speed-accuracy tradeoff. Finally, computational modeling indicated that motion percepts are accounted better by a Bayesian model in which Euler transformations are informed by priors on proprioception to estimate touch in body-centered coordinates. Collectively, these findings provide evidence for a flexible brain mechanism that maps tactile signals into task-relevant coordinate systems, with body-centered representations showing faster behavioral access.

## Methods

### Participants

A total of thirteen humans participated in Experiment 1 (∼300 total trials per participant). One participant was removed due to a lack of performance and non-attentiveness during the experiment, resulting in twelve participants (eight females). Eight participants were right handed, three ambidextrous, and one left handed, as assessed by the Edinburgh Handedness Inventory^51^. Experiment 1 was completed in one day of testing. A total of sixteen humans participated in Experiment 2. Two participants were removed due to a lack of comprehension of the task, as confirmed by participants themselves, resulting in a total of fourteen participants (eight females, all right-handed). All participants in Experiment 2 were unique from those in Experiment 1. Experiment 2 required a significant number of trials per participant (>2000 trials), completing the study across 4 to 5 days that resulted in 2300 - 2600 trials per participant.

The protocols for the experiments were approved by the Research Subjects Review Board of the University of Rochester, and written informed consent was obtained from all participants. Participants were remunerated for their participation in the experiment.

### Tactile Stimulator

We used a custom-designed 3D-printed cylindrical and dotted-pattern stimulus (32mm diameter) to provide tactile motion stimulation (**Figure 1A**). The dots were half-spheres with a radius of 2mm, and 5mm minimum surface distance between the dots. The wheel was aligned over the center of the distal phalange of digit 2 (D2, index finger) of the left hand. During alignment, we used a camera (Raspberry Pi Camera Module 3) to ensure that the wheel did not run over the edges of the finger. The wheel spun at 85 mm/s in one of twelve directions between 0º to 330° (steps of 30°) for a duration of 2 seconds. The stimulus was indented 1.5 mm into the skin surface.

**Figure 1.**
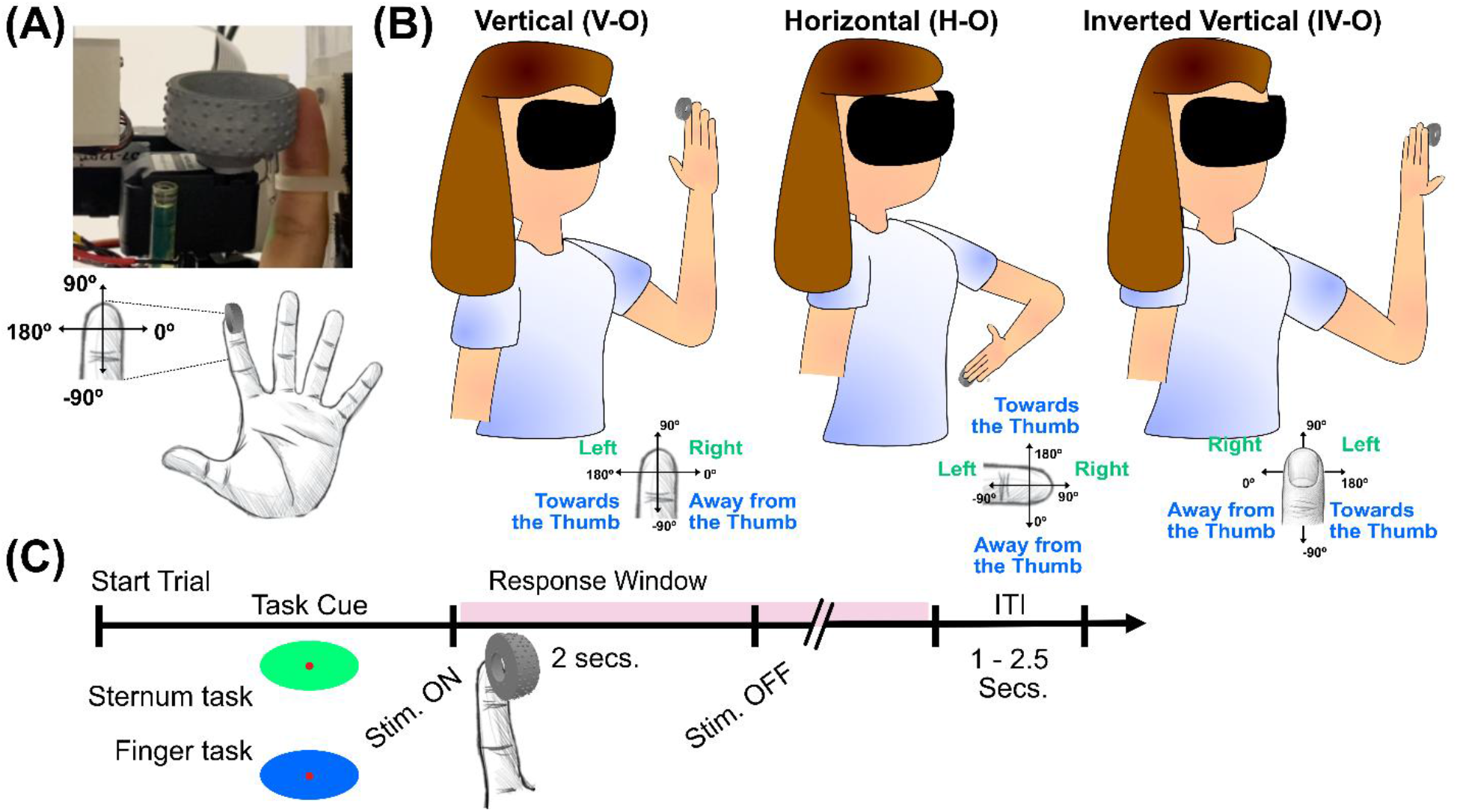
**(A) Tactile motion stimulator:** The top panel shows the dot-patterned cylindrical stimulus indented on participants’ index finger. The lower panel shows the angle directions of the tactile stimuli in relation to the anatomy of the finger (e.g., 90º is a motion stimulus moving towards the fingertip). **(B) Proprioceptive and reference frame task conditions:** The three proprioceptive conditions used in Experiment 1: Vertical-oriented (V-O, green left panel), Horizontal-oriented (H-O, blue middle panel) and Inverted Vertical-oriented (I-VO, red right panel). The insets in each condition show the discrimination axes for each reference frame task condition. The text in green and blue indicate the ideal responses in the Sternum- and Finger-centric tasks, respectively. **(C) Sequence of events in a trial:** Experimental timeline of a typical trial in Experiments 1 and 2.

### Experimental Setup and Task Design

The stimulated hand was secured in a custom-made exoskeleton using Velcro straps and finger splints to prevent finger movements. The nail of the stimulated finger was glued down to ensure that tactile stimulation was consistently presented on the same location and depth across trials. Participants’ heads rested on a chin rest device to limit head movements.

In Experiment 1, each participant sat in front of a table with their arm placed in one of three postures: Horizontally-oriented (H-O), Vertical-oriented (V-O), and Inverted Vertical-oriented (IV-O, **Figure 1B**). The forearm and elbow were rested for comfort in each of the three postures. Trials were blocked by posture in a randomized order. Within a block, participants were cued to judge the motion of tactile stimulation relative to the sternum (Sternum-centric task) or the thumb (Finger-centric task). In the Sternum-centric task participants determined whether the tactile stimulus moved towards the left or right in relation to the center of their body. In the Finger-centric task, participants indicated whether stimuli moved towards or away from the thumb on the stimulated hand. The two tasks were randomly interleaved within posture blocks, with a switching probability of 0.3 between tasks. On average, participants performed 5 trials per condition, resulting in a total of 120 trials per block. Participants completed one block of trials per posture condition. The total time of the experiment was between 1.5 and 2 hours.

The Sternum-centric task is akin to making judgments in body-centered coordinates, as it requires processing motion signals on the finger based on the position of the hand relative to the trunk of the body. For example, a stimulus moving at 0º on the finger with the hand in the V-O and IV-O postures should be judged moving rightward and leftward, respectively. The same motion stimulus should be perceived around the horizontal midline in the H-O posture, leading to equal leftward and rightward responses in the H-O posture. The Finger-centric task is akin to making judgments in skin-centered coordinates, as the same stimulus on the skin is perceived moving in the same direction regardless of hand posture condition. For example, a stimulus moving at 0º on the finger should be perceived as moving away from the thumb in the V-O, IV-O, and H-O posture conditions.

At the beginning of the experiment, participants performed a set of test trials to get accustomed to the task. Participants performed the experiment wearing a Virtual reality headset (Rift S, Oculus) to block visual inputs related to the hand position or tactile stimulation. **Figure 1C** shows the timeline of events for the experiment. A trial began with the presentation of a visual stimulus that cued the reference frame task condition (Finger-centric cue = blue oval, Sternum-centric cue = green oval; 500ms duration). Afterwards, the tactile stimulus was presented. Participants were instructed to make a response as soon as they felt confident in their decision. Five participants made responses via mouse clicks. In the Sternum-centric task, a left or right button press indicated that the stimulus moved leftward or rightward, respectively. In the Finger-centric task, a left or right button press indicated that the stimulus moved towards or away from the thumb, respectively. Although responding with button presses (mouse or foot pedals) is the traditional method employed in these types of discrimination tasks^6,20,38,52–58^, there are drawbacks in using button clicks to report behavior. In particular, tasks that require the same button press to report the perceptual decision of two or more stimulus conditions have the limitation of not knowing, in an unequivocal sense, whether incorrect responses were a result of an error in the perceptual decision itself or a lapse in the cued task that caused participants to judge stimuli in the incorrect reference frame. Further, button responses can induce cognitive mapping effects between button locations and discrimination spaces that may modulate RTs in a particular reference frame task^59,60^ (e.g., associating left button clicks to leftwards decisions might be easier to establish than left button clicks to ‘moving towards the thumb’ decisions). Given these limitations, the remaining seven participants performed the task by making verbal responses. Importantly, while participants were significantly faster in responding with the mouse as compared to verbally (F(1, 67) = 14.744, *p = 2.5 × 10*^*-4*^), there were no RT differences of hand posture angle (F(2, 67) = 1.3421, p = 0.27) or reference frame task conditions between the two modes of responding (F(1, 67) = 1.7928, p = 0.18; see **Supplementary Figure 1**). As such, responses were collapsed across these two response type conditions. We note that verbal responses have been used in other tactile discrimination tasks^9,61^.

The timeline of events in Experiment 2 was the same as in Experiment 1 (**Figure 1C**). Participants also performed the task under different proprioceptive states and reference frames wearing a Virtual reality headset (HTC Vive Pro, Microsoft). The proprioceptive state of the arm was changed randomly on every trial between five positions (15º, 25º, 30º, 50º, and 80º four participants did not undergo the 30º posture angle). Participants were cued to judge motion relative to the sternum (Sternum-centric) or the index finger on the hand (i.e., the stimulated finger, Finger-centric task), but the response axis was changed in the Finger-centric task (as compared to Experiment 1) to test whether our model is generalizable to other reporting coordinates. Specifically, participants judged the direction of motion of stimuli ‘left’ or ‘right’ relative to the sternum (Sternum-centric task) or ‘towards’ vs. ‘away from ‘the base’ of the stimulated finger (Finger-centric task). Responses in Experiment 2 were made verbally.

### Behavioral data analysis

Psychometric curves were plotted as a function of motion direction on the skin. For each participant, we transformed the psychometric function of each posture and task condition into a vector distribution. The length of each vector equals the proportion value for that angle vector in the psychometric curve. We then computed the circular mean of each vector distribution to statistically test for posture-dependent shifts in participants’ response distributions (see **Supplementary Figure 2**). We performed Watson-Williams one-way ANOVAs for circular data to tests for posture-dependent shifts in the mean distributions across proprioceptive conditions. Participants’ reaction times (RTs) were calculated by averaging their RT across all trials within each reference frame and posture angle condition. Statistical testing of the RT data was performed by submitting the data to generalized linear models (GLMs), using an inverse Gaussian function. Pairwise Ranksum tests were conducted to test for differences between conditions.

To formalize how proprioceptive and tactile signals are flexibly weighted might be implemented, we introduce a simple task-weighted model in which the estimated direction of motion is expressed as a linear combination of skin-based motion signals and posture-dependent transformations of those signals (**Equation 1**):

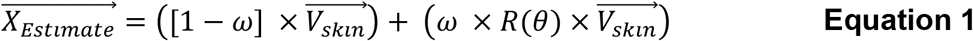

where 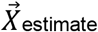 is the estimated direction of motion, 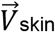 is the direction of motion of a stimulus in skin coordinates, R is a transformation applied on 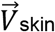 based on the posture angle θ to represent touch in non-skin coordinates, and ω is the weight value applied to the sensory signals (which can be task-dependent). The strategy for an ideal observer is to set ω = 0 for tasks that do not require proprioception, and ω = 1 for tasks that require remapping of skin signals into a coordinate frame anchored to the sternum.

### Computational modeling

#### Accumulation to bound model

To determine whether differences in RTs across reference frame conditions reflect changes in sensory evidence accumulation, decision policy, or both, we applied a drift diffusion model (DDM) to participants’ trial-by-trial choices and RTs adapting code from a publicly-available DDM software package^62,63^. The DDM provides a mechanistic account of two-alternative decisions by modeling behavior as a stochastic accumulation of noisy sensory evidence toward one of two discrimination boundaries^21,22,64^. Formally, the decision variable x(t) evolves according to the stochastic differential equation (**Equation 2**).

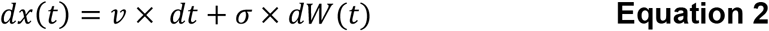

where ν denotes the drift rate (i.e., mean rate of evidence accumulation), σ denotes the noise amplitude (fixed to 1 for identifiability), and dW(t) represents a Wiener noise process. We used the standard 0 → ‘a’ formulation of the DDM, in which the decision variable evolves between a lower bound at 0 and an upper bound at ‘a’, where ‘a’ denotes the decision threshold estimated from the data. The starting point (z) was expressed as a fraction of the boundary separation and was fixed at the midpoint (z = a / 2), reflecting a symmetric prior over the two response alternatives within each task. Decision threshold and starting point varied by reference frame task. A decision is registered when x(t) reaches either boundary, which determines the choice. RT is given by the sum of the decision time and the non-decision time component (T_ER_), which estimates brain processes not directly related to evidence accumulation. T_ER_ was estimated by pooling data across hand posture conditions.

Sensory evidence was defined on each trial as the signed angular distance between the stimulus direction and the task-relevant discrimination boundary. In the Finger-centric task, the discrimination boundary was fixed across postures. In the Sternum-centric task, the boundary was rotated according to hand posture, such that evidence was computed relative to a posture-dependent boundary on each trial. For example, if the base Sternum-centric boundary is 90° (i.e., motion along the long axis of the finger in the vertical-oriented posture), and the elbow is rotated by +90° about the z-axis (horizontal-oriented posture), the effective decision boundary rotates to 180°. Thus, a stimulus moving with an angular direction of 170° would then lie 10° away from the discrimination boundary, whereas the same stimulus in the vertical posture would lie 80° away from the discrimination boundary. After this transformation, trials were pooled across postures, and the DDM was fit with task-specific drift gains and decision thresholds, and a shared non-decision time (Ter). Pooling the posture-transformed data allows us to isolate task-level differences in decision dynamics while ensuring sufficient trial counts for stable parameter estimation and statistical power. Signed evidence for trial ‘i’ (e_i_) was computed using **Equation 3**

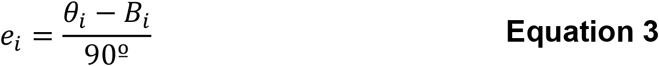

where θ denotes stimulus angle and B the corresponding discrimination boundary angle. The denominator of 90° reflects the maximum possible signed angular distance to a discrimination boundary in a two-alternative circular task (±90°), effectively normalizing signed evidence between ±1. For example, a stimulus 45° from the boundary yields e = 45º / 90º = 0.5, whereas a stimulus orthogonal to the boundary (i.e., 90° away) yields e = 1. Angular differences were computed on the circle by wrapping angles to the range ±180°, and calculating the signed distance to the nearest of the two opposing discrimination boundaries (B and B + 180°). As a result, angular deviations larger than 90° were automatically mapped to the opposite boundary, ensuring that evidence magnitude did not exceed ±1 after normalization. Evidence was signed such that positive values favored the response coded as 1 (e.g., toward the thumb in the Finger-centric task). Additional task-specific boundary offset parameters (δ_sternum_, δ_finger_) were included to capture systematic shifts in categorical decision criteria.

Drift rate (ν) was modeled as proportional to trial-wise evidence, with separate gain parameters for each reference frame (see **Equation 4**)

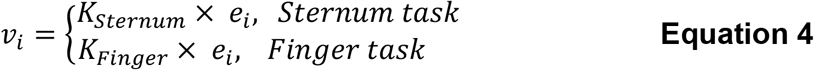

where K_sternum_ and K_Finger_ represent the sensitivity in each task, and were constrained to be larger than 0. Decision thresholds were allowed to vary by task (a_Finger Task_ and a_Sternum task_). Larger decision thresholds (or boundary separation) reflect a more conservative decision criterion (i.e., requiring more accumulated evidence before committing to a response). Response bias was modeled as a task-specific starting-point bias separately (‘z’), allowing the initial value of the decision variable to deviate from the midpoint between bounds and thereby capturing an a priori preference for one response alternative. An additive drift bias term was not included in the model.

Because drift rate (ν) varied across trials as the product of signed evidence (e) and an evidence accumulation gain term (K), we report the task-specific gain parameter (K) rather than mean drift rate. This parameter more directly reflects the sensitivity of the accumulation process to sensory evidence (see Materials and Methods). Critically, stimulus directions were sampled symmetrically around the discrimination boundary, yielding signed evidence values centered near zero, with slight negative offsets in some participants. As a result, average drift rates can appear small or even negative due to the stimulus distribution rather than weak evidence accumulation per se. Reporting K therefore provides a more interpretable measure of accumulation sensitivity, independent of the geometry of the stimulus set. Although drift gain (K) and decision threshold (a) are often interpreted independently, their effects on decision time (T_dec_) are intrinsically nonlinear. For constant drift rate (ν), the expected decision time (excluding T_ER_) is approximated by **Equation 5** ^24–26^:

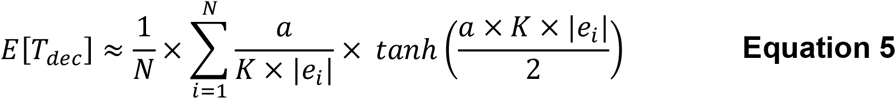

where N is the total number of trials. The key component for estimating E[T_dec_] is to take the absolute value of the signed evidence (e) on every trial. **Equation 5** illustrates that RT does not necessarily scale linearly with either drift rate or decision threshold alone. Consequently, a reduction in decision threshold can yield faster RTs even when drift (or drift gain) is smaller, because the mapping from (v and/or a) to T_dec_ is nonlinear.

Parameters for the DDM model were estimated at the individual participant level using maximum likelihood estimation. Optimization was initialized from multiple random starting points to ensure robust convergence, and the parameter set yielding the lowest negative log-likelihood was retained. To assess parameter reliability and support statistical inference, nonparametric bootstrapping was performed by resampling trials with replacement within each participant (n = 1000). Confidence intervals on task-dependent parameter differences were computed from the resulting bootstrap distributions.

#### Coordinate transformation modelling

To model how tactile motion signals are transformed from a Finger-centric (i.e., skin-based) to a Sternum-centric (i.e., body-centered) reference frame, we used Euler rotations as a compact and principled framework for mapping vectors between coordinate systems. Euler transformations provide a minimal parameterization of posture-dependent rotations, naturally capturing the hierarchical structure of the upper limb, in which rotations at proximal joints propagate to distal segments. In our task, proprioceptive-dependent transformations were implemented as Euler matrix rotations about the elbow joint (z-axis rotation), allowing skin-based motion vectors to be systematically remapped into Sternum-centered coordinates based on arm posture **(Equation 6)**.

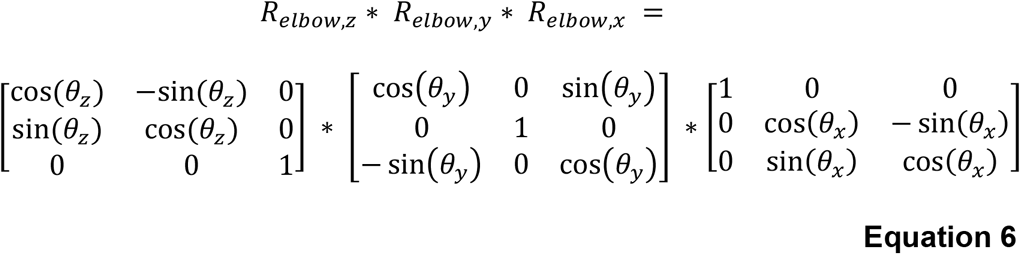

where R_elbow,x_, R_elbow,y_, R_elbow,z_ refer to the rotation angle of a body part around the x, y, and z axes. θ equals the brain representation of v_p,_ the transduced proprioceptive state of a body part in x, y, and z coordinates. Rotating the elbow from 15º to 80º, as in Experiment 2, requires elbow rotations around the Z-axis only (**Equation 7**).

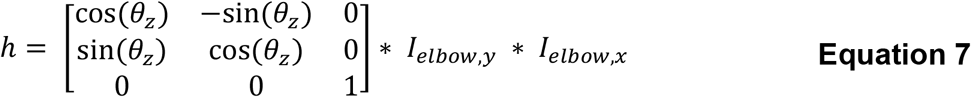

Here, I_x_ and I_y_ are identity matrices around the X and Y axes, and *h* represents the Euler transform. Our model makes a choice (e.g., left or right relative to the sternum) based on the reference frame condition given a physical stimulus, *e*_*m*_, represented in the brain as S_Finger_. The transformation from S_Finger_ to S_Sternum_ can be approximated by **Equation 8**.

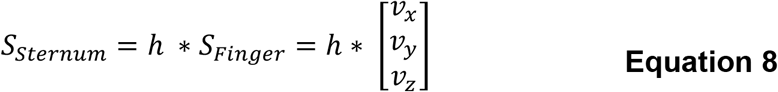

where the vector (v_x_, v_y_, v_z_) denotes the representation of the direction of motion of the stimulus in the brain (S_Finger_) that can be estimated by the Full Vector Average model^28,29^ (see supplementary material for more details underlying the Euler model). The Finger-and Sternum-centric tasks were fit separately since the weight model analysis (**Equation 1**) indicated largely independent effects of tactile and proprioceptive signals across the two reference frame task conditions, with slopes near the theoretical predictions for each frame. This separation suggests that participants largely adopted the instructed reference frame in each task, allowing the Bayesian model to be applied independently for the two conditions.

We developed a Bayesian model that allows for nonuniform *priors* over arm posture and choice in each task. Derivations of the Euler and Bayesian models are described in detail in the supplementary material. For all models, we minimize the negative log-likelihood of the parameters to get the best fit of the model using the BADS algorithm^65^ implemented in MATLAB (R2021a, Mathworks Inc.). The optimization procedure was repeated 20 times with different initial parameters. The final parameters were largely insensitive to initial values. We use the likelihood-based pseudo-R^2^ measure^66,67^ (**Equation 9)**:

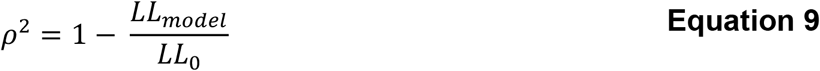

where LL_model_ is the log-likelihood of the model and LL_0_ is the log-likelihood of the null model, which only includes an intercept as the predictor. In general, the values corresponding to a good fit are lower for ρ^2^ as compared to R^2^, with ρ^2^ values ranging from 0.2 to 0.4 considered to be excellent fits to the data^66,67^.

## Results

### Tactile and proprioceptive signals are combined based on task demands

Participants flexibly discriminate tactile motion stimuli in different reference frames. **Figure 2A** shows largely overlapping vs. divergent psychometric curves across hand posture conditions for stimuli judged relative to the finger (*left panel*) vs. the sternum (*right panel*), respectively. Watson-Williams multi-sample tests showed a shift in circular mean across hand posture angle conditions for the Sternum-centric (F(2,33) = 260.42, p < 10^−15^), but not the Finger-centric task (F(2,33) = 0.46, p = 0.63; see **Supplementary Figure 2**). Post-hoc Watson-Williams uni-sample tests in the Sternum-centric task revealed significant differences between the V-O vs. H-O postures (F(1,22) = 191.23, *p = 2.49×10*^*-12*^, Δ circular mean = 118º), V-O vs. IV-O postures (F(1,22) = 1.7199 × 10^3^, p < 10^−15^, Δ circular mean = 177º), and IV-O vs. H-O postures (F(1,22) = 47.29, p < 6.60 × 10^−7^, Δ circular mean = 58º). **Supplementary Figure 3** shows participants’ responses plotted as a function of the motion direction in coordinates relative to the sternum, with psychometric curves converging across hand postures in the Sternum-centric task and diverging in the Finger-centric task. This pattern reflects reference frame-dependent remapping, indicating that participants evaluated the direction of motion in the task-relevant coordinate system. Taken together, these data provide empirical evidence that reference frame signals control how tactile and proprioceptive inputs are integrated to generate touch perception in the task-relevant reference frame.

**Figure 2.**
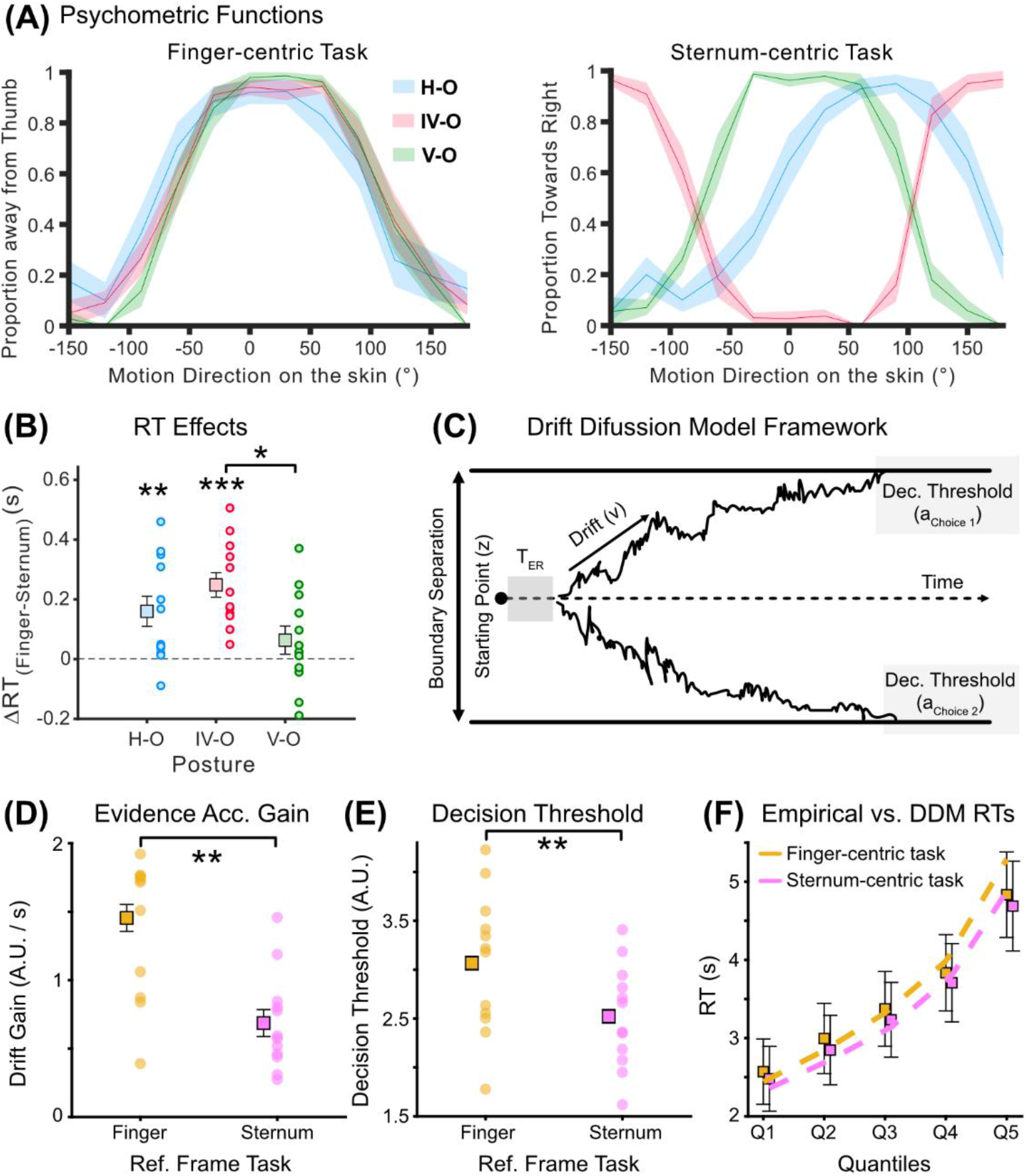
**(A) Motion discrimination results for Experiment 1:** Group-averaged psychometric functions in the Finger-centric (left) and Sternum-centric (right) tasks for each posture angle condition. Shaded error bars represent standard error of the mean (SEM) across participants. **(B) Reaction Time (RT) effects across reference frames:** Difference in RTs between the Finger- and Sternum-centric tasks for each posture angle condition (H-O = blue, IV-O = red, and V-O = green circles). The square objects represent the mean ΔRT for each posture angle. **(C) Schematic of the drift diffusion model (DDM):** Noisy sensory evidence is accumulated over time from a starting point until it reaches one of two decision thresholds, determining choice and reaction time. The model parameters include drift rate (‘v’), decision threshold (‘a’), starting point (‘z’), and non-decision time (T_ER_). Boundary separation reflects the distance between the decision thresholds for both choices. **(D) Evidence accumulation across tasks:** Drift rate estimates from the DDM plotted separately for Finger- and Sternum-centric tasks. The square objects represent the mean drift rate for each task. **(E) Response boundaries across tasks:** Decision threshold estimates from the DDM plotted separately for Finger-and Sternum-centric tasks. Conventions as in (D). **(F) DDM fit to empirical data:** Quantile–quantile comparison of empirical (boxes with error bars) and DDM-simulated RTs (dashed lines) for each task condition. Error bars represent within-subject SEM as computed by ^68^. N = 12.

**Figure 3.**
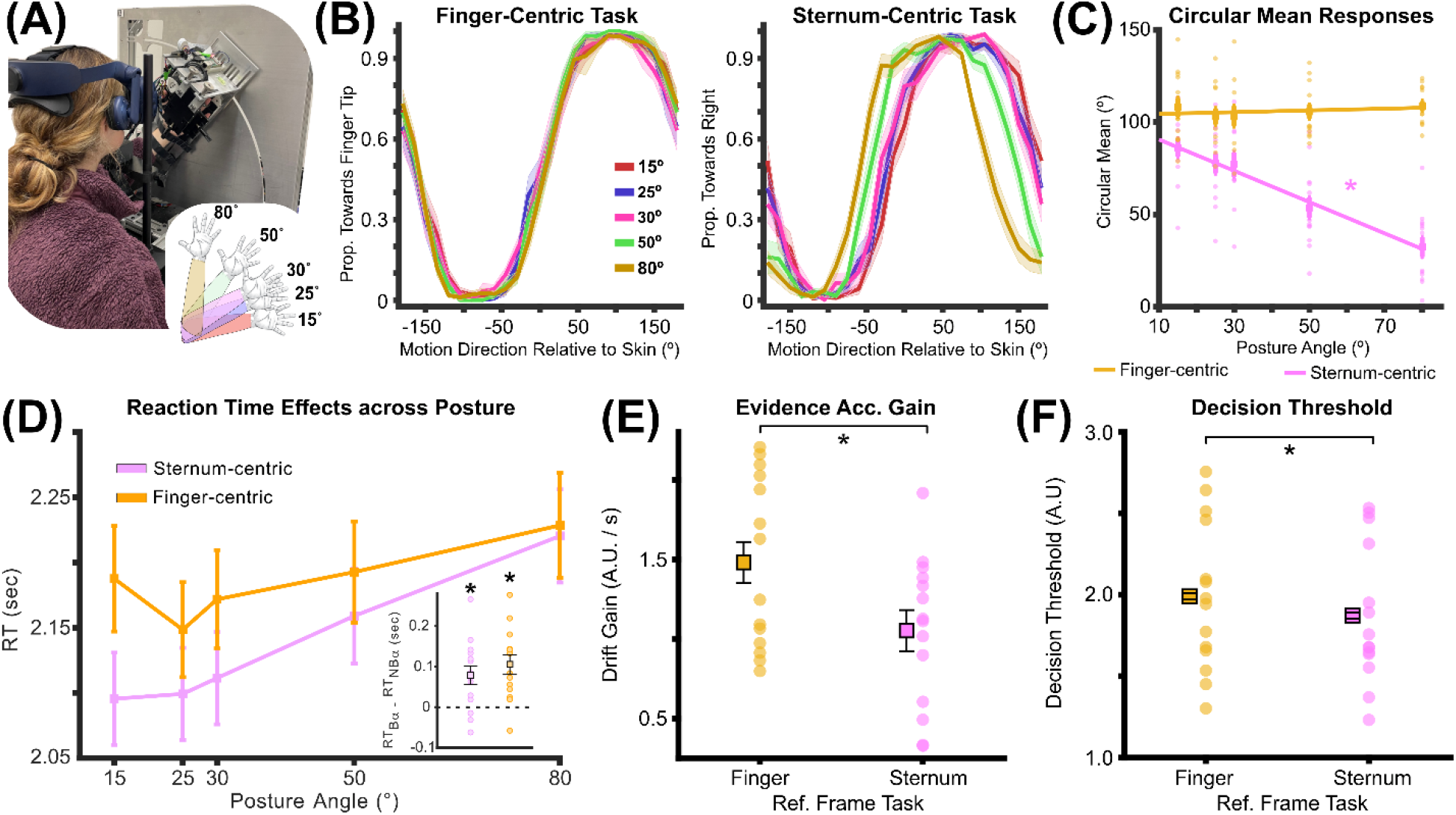
**(A) Experimental setup for Experiment 2:** Picture of the experimental device used to systematically modulate the posture of the arm, and present tactile motion stimuli to the index finger. The inset represents the different posture angles used in Experiment 2. **(B) Motion discrimination results for Experiment 2:** Psychometric functions averaged across participants in the Finger-centric (left) and Sternum-centric (right) tasks for each posture condition in Experiment 2. Shaded error bars represent SEM across participants. Similar to Experiment 1 data, there is a systematic shift in the psychometric functions when participants are performing the Sternum-centric task only. **(C) Circular mean of psychometric curves:** Circular means across participants as a function of posture angle for the Sternum-centric task (magenta trace) and Finger-centric task (orange trace). **(D) Reference frame and posture effects on RTs:** RT data across posture angles for the Finger-centric (orange trace) and Sternum-centric (magenta trace) task conditions. The insets are averaged RT difference between stimuli moving in the NBα and Bα conditions. **(E) Evidence accumulation across tasks in Experiment 2:** Drift gain estimates from the DDM plotted separately for Finger- and Sternum-centric tasks. The square objects represent the mean drift rate for each task. **(F) Response boundaries across tasks in Experiment 2:** Decision threshold estimates from the DDM plotted separately for Finger- and Sternum-centric tasks. Conventions as in (E). Error bars represent within-subject SEM as computed by ^68^. N = 14.

### Motion judgements in body-centric coordinates are faster and driven by lower decision thresholds

Response times (RTs) are faster in the Sternum-centric task. **Figure 2B** shows differences in RTs between reference frame tasks for each hand posture angle condition. A generalized linear model (GLM) revealed a main effect of reference frame task condition on RT (F(1, 22) = 18.95, *p = 2.54 × 10*^*-4*^), with faster RTs for the Sternum-(mean RT = 2.92 seconds) relative to Finger-centric task (mean RT = 3.04 seconds). The GLM did not show a main effect of hand posture angle (F(2, 33) = 0.78, p = 0.466), but did reveal an interaction effect between reference frame task and hand posture angle (F(2, 66) = 4.33, *p = 3.82 × 10*^*-3*^). Post-hoc Wilcoxon signed-rank tests revealed faster RTs in the Sternum-vs. Finger-centric condition for the H-O (Z = 3.67, *p = 2.41 × 10*^*-4*^) and IV-O (Z = 4.41, *p = 1.03 × 10*^*-5*^) postures, but not for the V-O posture (Z = 1.45, p = 0.14). The analysis also showed greater differences between the Finger- and Sternum-centric tasks for IV-O vs. V-O posture conditions (Z = 2.57, *p = 1.02 × 10*^*-2*^). No other significant differences were observed. Together, these results show that Sternum-centric judgments are executed more rapidly than Finger-centric judgments, suggesting that information represented in body-centered coordinates is accessed more rapidly during decision making.

Angles near the discrimination boundary were slower to discriminate in both reference frame tasks (**Supplementary Figure 4**). Importantly, the angles associated with slower RTs shifted across hand postures in the Sternum-centric task, reflecting the expected posture-dependent shifts in the discrimination boundary, but remained invariant across hand posture in the Finger-centric task. Together, these results further suggest that participants performed motion discrimination in the instructed reference frame, as RTs peaked near the reference frame-specific discrimination boundary and only shifted with posture in the Sternum-centric task.

**Figure 4.**
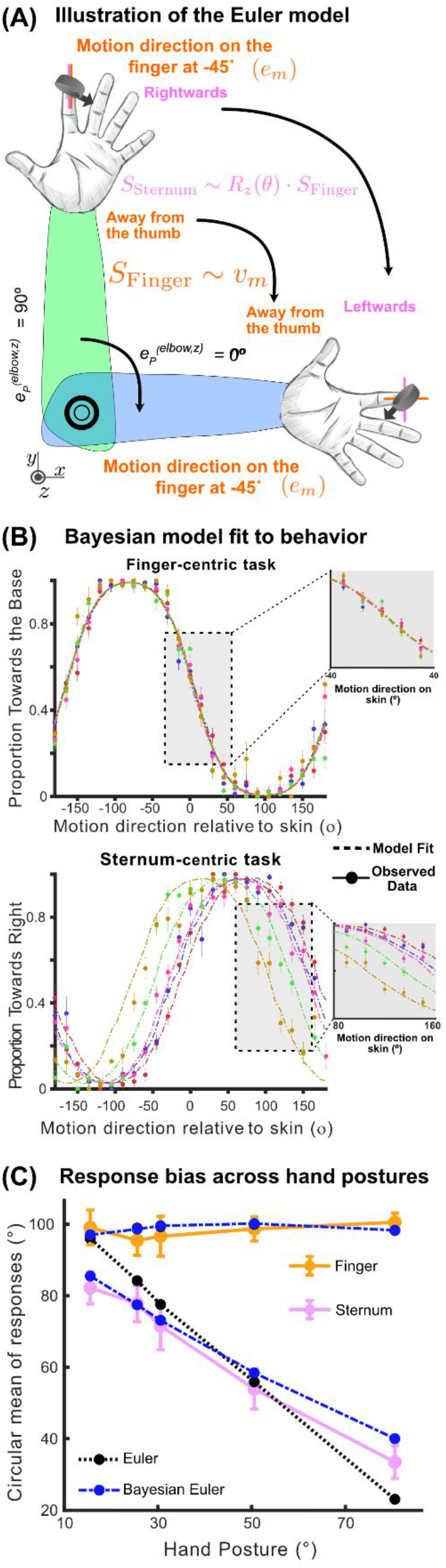
**(A) Euler model example and predictions:** The forearm moving from a Vertical-oriented (V-O, *green*) to Horizontal-oriented (H-O, *blue*) posture, and the Euler model’s response to a stimulus moving at -45° (e_m_) made in a Finger-centric and Sternum-centric reference frame. In the V-O posture (elbow posture angle = 90º; e_p_), the ideal subject responds ‘rightward’ in the Sternum-centric task and ‘away from the thumb’ in the Finger-centric task. The arm is rotated around the elbow by 90° along the z-axis to establish an H-O posture (elbow posture angle 0º). In this new posture, the ideal subject responds leftwards in the Sternum-centric task to a stimulus moving at -45º and away from the thumb in the Finger-centric task. v_m_ is the neural sensory representation of e_m_. S_Finger_ represents the perceived direction of motion in the Finger-centric reference frame, which is approximated by v_m_ (as predicted by the Full Vector Average model^29^). S_Sternum_ represents the perceived direction of motion in the Sternum-centric reference frame, computed via an Euler transform (R_z_(θ)) of v_m_. **(B) Bayesian model fits to the data in Experiment 2:** Bayesian model fits to the psychometric functions in **Figure 3B** for the Finger-centric (top) and Sternum-centric (bottom) task in each posture angle condition. **(C) Response bias across hand postures for modeled and empirical data:** Mean circular response angle plotted as a function of hand posture for the Finger-centric (orange) and Sternum-centric (magenta) tasks. Dashed and dotted lines show model predictions derived from the Euler (black, dotted) and Bayesian model (blue, dashed). The Bayesian model best estimates the systematic posture-dependent shifts in circular mean observed in the Sternum-centric condition.

Prior work has shown that visual motion direction discrimination is well described by evidence-accumulation models, in which both sensory evidence and decision policy jointly shape choice and RT^21–23^. We applied a drift diffusion model (DDM) to determine whether faster RTs in the Sternum-centric task reflect differences in drift rate, decision thresholds, or both (**Figure 2C**). The DDM was fit to each participant’s trial-by-trial choice and RT data, jointly modeling Finger- and Sternum-centric trials within a single framework (see Methods for details). Because drift rate varied across trials as the product of signed evidence (‘e’) and the evidence accumulation gain term (‘K’), we report the task-specific evidence accumulation gain parameter (as opposed to drift rate), which more directly reflects accumulation sensitivity (see Materials and Methods).

A Wilcoxon signed-rank test revealed higher drift gain in the Finger-vs. Sternum-centric task (1.45 vs. 0.69; Z = 3.05, *p = 0.002*), indicating stronger sensory evidence accumulation in the Finger-centric condition (**Figure 2D**). However, we observed lower decision thresholds for the Sternum -centric task (3.07 vs. 2.52; Z = 2.98, *p = 0.003*), suggesting that participants needed less time to accumulate evidence in the Sternum-centric condition before committing to a response (**Figure 2E**). A Wilcoxon signed-rank test against 0.5 did not reveal differences in the starting point of evidence accumulation for the Finger-centric task (Z = -1.09, p = 0.27) or Sternum-centric task (Z = 0.54; p = 0.58). Although drift gain is lower in the Sternum-centric task, RTs are faster. This apparent mismatch likely reflects a fundamental property of the DDM, wherein mean decision time depends nonlinearly on both drift rate parameters and decision thresholds^24–26^. Specifically, under symmetric bounds and constant drift, the analytical solution of the expected decision time follows a hyperbolic tangent-shaped function of the interaction between drift and decision threshold distance between the two choices (i.e., boundary separation) that reflects the time for accumulated evidence to reach a bound of the Wiener diffusion process (**Equation 5** in Methods)^25^. As such, RT does not scale linearly with drift gain alone. In the fitted parameter regime, the reduction in decision threshold in the Sternum-centric task shortens the distance to bound sufficiently to offset its lower drift gain. Consistent with this hypothesis, we found that the estimated decision time computed using **Equation 5** was strongly correlated with participants’ empirical RTs for the Finger-centric task (R = 0.85, p = 4.43 × 10^−4^) and the Sternum-centric task (R = 0.84, p = 5.81 × 10^−4^; **Supplementary Figure 5**).

We compared empirical and simulated RTs using quantile plots to evaluate correspondence across the RT distribution for each participant. The DDM closely reproduced the shape of the RT distribution in both tasks, with systematically increasing RTs across quantiles and consistently faster responses in the Sternum-centric condition (mean R^2^ across participants = 0.60 and 0.83 for the Finger- and Sternum-centric tasks, respectively; **Figure 2F**). Importantly, the quantile analysis shows that the faster RTs in Sternum-centric is present across the entire RT distribution, indicating that the RT benefit reflects a global shift in decision time rather than a selective difference in a subset of trials. To further demonstrate that the faster RTs in the Sternum-centric task reflect lower decision thresholds, we refit the DDM to the empirical data while systematically swapping drift gain and/or decision threshold parameters across tasks to test whether the resulting parameter combinations reproduce the empirical RT patterns. If the RT advantage arises from lower decision thresholds in the Sternum-centric task, then swapping the decision thresholds across tasks should reverse the RT ordering, whereas swapping drift gain alone should not reproduce the empirical RT difference. Consistent with this prediction, swapping the decision thresholds reversed the Sternum-centric RT advantage (**Supplementary Figure 6A**), whereas swapping drift gain produced RT predictions that fell well outside the empirical variability (**Supplementary Figure 6B**). Swapping both parameters yielded a combination of these effects (**Supplementary Figure 6C**). Collectively, these analyses indicate that the Sternum-centric RT advantage is primarily driven by reduced decision thresholds rather than increased evidence gain, consistent with a more liberal decision policy that resemble a speed-accuracy tradeoff.

### Tactile motion perception reflects flexible weighting of tactile and proprioceptive cues

Although Experiment 1 demonstrates that tactile and proprioceptive signals are not combined with fixed weights across reference frames (**Figure 2A**), it remains unclear how these signals are weighted to generate perceptual reports in the task-relevant reference frame. In Experiment 1, posture manipulations were not parametrically varied along a defined rotational axis, making it difficult to determine whether task-dependent weighting follows the principled transformation specified in **Equation 1**. As such, in Experiment 2, we implemented an experimental design in which the elbow was systematically rotated about the z-axis (from 15° to 80°). Participants discriminated tactile motion on the index finger in a Finger-centric (towards vs. away from the fingertip) or a Sternum-centric reference frame (left vs. right; **Figure 3A**). This experimental setup enables us to assess whether perceptual reports vary systematically with arm posture based on the task-relevant reference frame. Under **Equation 1**, the slope of this relationship provides a behavioral readout of the effective transformation of tactile signals across postures that is indexed by the weight (ω). Larger weight values indicate greater incorporation of posture information.

Similar to Experiment 1, Watson-Williams tests showed that hand posture angle modulates the circular means of motion judgements in Sternum-centric (F(4,61) = 14.11, *p = 3.27 × 10*^*-8*^) but not Finger-centric reference frames (F(4,61) = 0.21, p = .93; **Figure 3B**). A post-hoc linear regression analysis showed a systematic decrease in circular mean as a function of hand posture angle for the Sternum-centric task (F(1,64) = 57.26, *p = 1.87 × 10*^*-10*^, slope = -0.85; **Figure 3C**). These behavioral data align with a task-dependent weighted integration model. In the ideal case, Finger- and Sternum-centric judgments would correspond to slope values of 0 and −1, respectively. While the Finger-centric slope matches this prediction, the Sternum-centric slope was −0.85, reflecting a slight deviation from the ideal weight value that may arise from posture-dependent biases^27^.

### Reaction times are modulated by proprioception in a reference frame-dependent manner

Similar to data in Experiment 1, a GLM test revealed slower RTs in the Finger-centric vs. Sternum-centric condition (F(1,128) = 10.80, *p = 0.001;* **Figure 3D**). The GLM also revealed an effect of hand posture angle on RTs (F(1,128) = 23.72, *p = 3.22 × 10*^*-6*^). Follow up tests, using regression analyses, revealed a positive linear relationship between RT and Postural Angle condition in the Sternum-centric (F(1,67) = 27.29, *p = 1.8 × 10*^*-6*^; R^2^ = 0.29), and a weaker relationship for the Finger-centric reference frame task condition (F(1, 67) = 4.83, *p = 0.03*, R^2^ = 0.06). The GLM on the RT also showed a trend towards a significant interaction effect between reference frame task and hand posture angle (F(1,128) = 3.43, p = 0.066). Finally, Wilcoxon signed-rank tests also revealed higher RTs for angles near the discrimination boundary (*B*α) vs. those away from the discrimination boundary (*NB*α) for the Finger-centric (Z = 4.10, *p = 4.12 × 10*^*-5*^) and Sternum-centric condition (Z = 2.73; *p = 6.4 × 10*^*-2*^; see inset **Figure 3D**). Taken together, these data provide further evidence that perceptual judgements of tactile motion are faster when made relative to the sternum.

Similar to the data in Experiment 1, DDM findings revealed higher drift gain in the Finger-vs. Sternum-centric task (Wilcoxon signed-rank test; Z = 2.35, *p = 0.0186;* **Figure 3E**). We also observed lower decision thresholds for the Finger-vs. Sternum-centric task (Z = 2.61, *p = 0.009*; **Figure 3F**). However, unlike Experiment 1, we found a difference in the starting point of evidence accumulation between in the Finger-centric task (Z = 2.17, *p = 0.03*), indicating a slight bias for ‘towards the fingertip’ (0.53 vs. 0.47). We did not observe a difference in starting point of evidence accumulation for the Sternum-centric task (Z = -0.28; p = 0.78). Collectively, these findings converge with those from Experiment 1 by showing that judgments made in a Sternum-centric reference frame are characterized by lower decision thresholds, consistent with a faster commitment in decision making.

### A Bayesian framework mediates flexible perception of tactile motion

The prevailing model of tactile motion, the full vector average (FVA) model, estimates the direction of motion in a skin-centered reference frame only^28,29^. As such, the FVA cannot account for the reference frame-dependent behavior observed here, indicating that this model needs to be extended to integrate tactile and proprioceptive signals in a task-dependent manner. A natural computational framework for performing transformations across multi-joint rotations comes from robotics and motor control, where Euler rotations provide a compact set of matrix operations that map vectors across coordinate systems. Here, we developed a model in which vectorized tactile motion signals on the skin are transformed by Euler rotations based on arm posture (**Figure 4A**). However, sensory representations and perceptual decisions are inherently noisy and subject to individual biases^30–36^. Some studies show that proprioceptive representations of limb position are stable, but idiosyncratic across individuals, suggesting that deviations from veridical transformations may reflect priors over one’s own arm posture^37^. Consistent with this view, recent work demonstrates that errors in hand localization emerge from Bayesian integration of noisy sensory signals^27^. In line with this account, we observed that, in the Sternum-centric task, participants’ responses across different hand postures deviate from the identity line (**Figure 3C**; slope = −0.85), consistent with a bias in hand position estimates. Further, our data reveal idiosyncratic choice biases across participants in both the Sternum- and Finger-centric tasks (**Supplementary Figure 7**). Some participants show a stronger tendency to report motion ‘towards the thumb’ in the Finger-centric task or ‘towards the right’ in the Sternum-centric task, whereas others exhibit the opposite pattern. Collectively, these observations motivate a Bayesian extension of the Euler-based transformation framework that incorporates individuals’ priors over posture angle and task-dependent decision variables (see Methods and Supplementary Materials).

We developed a Bayesian model that provides good fits to participants’ behavior across reference frames and proprioceptive conditions (**Figure 4B**, *dashed traces;* see Methods and Supplementary Material for details on the model), with robust performance at the individual-subject level (**Supplementary Figure 8**). To determine which model best describes the empirical data, we fitted each model to individual participants’ data from Experiment 2 and compared Akaike Information Criterion (AIC) values across models, with ΔAIC < −20 indicating strong preference for the Bayesian model over the Euler model. Under this criterion, the Bayesian model outperformed the Euler model for eleven out of the fourteen participants (**Supplementary Table 1**). Consistent with this pattern, the Bayesian model showed a tighter correspondence between predicted and empirical circular means across hand postures, indicating that incorporating priors on hand posture and choice effectively captures the systematic posture-dependent biases observed in behavior (**Figure 4C**). Within the framework of **Equation 1**, these results provide a mechanistic interpretation of the posture-dependent biases observed in behavior for the Sternum-centric task (**Figure 3C**). The deviation from the identity line indicates that posture information is not incorporated veridically, and the Bayesian model accounts for this bias through a prior. Under this formulation, the weight value (ω) in the Sternum-centric task likely reflects the transformation of tactile signals based on an inferred estimate of posture, shaped by both sensory evidence and priors, rather than interference between reference frame tasks. In sum, these findings indicate that while Euler-based transformations provide a principled framework for reference frame computations, incorporating Bayesian priors over arm posture and choice accounts for systematic under-correction of tactile motion discrimination across hand posture.

## Discussion

We investigated how proprioceptive signals and task demands shape the perception of tactile motion in different reference frames. Our results show that the brain flexibly integrates tactile and proprioceptive information in a task-dependent manner, with proprioception systematically modulating motion judgments only when participants judge motion relative to the sternum. These findings challenge prior accounts by showing that tactile motion reports are not uniformly biased by arm postural changes^6,20,38^. Specifically, Chen and colleagues^20^ reported that tactile motion judgments depend on the relative position of the finger and head, whereas other studies propose that proprioceptive inputs are automatically integrated with tactile signals^6^. In contrast, we found that tactile motion judgments made in a Finger-centric reference frame were invariant to hand posture (**Figures 2A and 3B**). These results indicate that integration of tactile and proprioceptive signals is flexible rather than obligatory, and depends on the reference frame required by the task. Consistent with this interpretation, the slope that associates perceived motion direction to posture angle is near zero for Finger-centric judgments, but approaches −1 for Sternum-centric judgments (**Figure 3C**), reflecting strong posture-dependent transformations when information needs to be reported in a body-centered frame. Together, these findings support a task-weighted scheme in which proprioceptive transformations are selectively engaged to map skin-centered motion signals into task-relevant coordinate systems, rather than a fixed or continuous integration mechanism across reference frames.

Analyses of the RT data reveal a consistent speed advantage for Sternum-centric over Finger-centric motion judgments (**Figures 2B and 3D**), an effect that replicated across both experiments. Accumulation-to-bound modeling shows that these faster responses are not driven by stronger or faster accumulation of sensory evidence, but by lower decision boundary thresholds. This dissociation is important because it indicates that faster Sternum-centric judgments do not necessarily reflect enhanced sensory precision, but rather a change in decision policy that favors earlier commitment. Such a strategy is well suited for dynamic interactions with objects, where it may be more critical for motor systems to access sufficiently reliable sensory signals with minimal delay, rather than fine-grained sensory detail. In this context, sacrificing precision for speed may be advantageous, as rapid access to body-centered motion information enables timely updates of posture-independent grasping in response to sudden grasp-failure events such as object slip. From this perspective, lower decision thresholds in the Sternum-centric task reflect a speed-accuracy tradeoff that prioritizes rapid sensorimotor readiness over maximal perceptual fidelity.

A Bayesian framework incorporating Euler matrix computations provides a robust account of how tactile motion signals on the skin are transformed by proprioceptive inputs in a reference frame-specific manner (**Figure 4**), and the inclusion of priors on hand posture yielded substantially better fits (**Table 1**). Euler-based transforms offer a principled advantage in their generalizability, particularly for humans and non-human primates who possess a large number of degrees of freedom in the upper limbs^39,40^ and joints arranged in a hierarchical and nested structure. Euler rotations provide a compact and scalable means of mapping touch across such multi-joint configurations without requiring task-or posture-specific reparameterization. These computations are also biologically plausible. Euler transformations can be expressed as combinations of sine and cosine terms^41,42^ that, theoretically, can be implemented through linear population readouts of neural ensembles. Consistent with this view, neurons in several sensory and motor cortical areas exhibit cosine-like tuning to joint angles^43–47^, providing a neural basis through which weighted combinations of proprioceptive signals may facilitate task-dependent transformations of sensory information in the brain.

A central question raised by our findings concerns the order in which tactile motion signals are represented across reference frames. Traditional models posit that tactile information is initially encoded in a skin-centered reference frame and subsequently transformed into body-centered representations in higher-order cortical areas^48,49^. More recent accounts, however, propose that tactile and proprioceptive signals are automatically integrated^6,38^, giving rise to body-centered representations at early stages of processing. Consistent with this latter view, we have shown that neurons throughout somatosensory cortex (including areas 3a and 3b) are responsive to both tactile and proprioceptive inputs^50^, indicating that circuits in early neocortex are well equipped to perform posture-dependent transformations. Our RT findings provide converging behavioral support for this hypothesis, in that Sternum-centric judgments are consistently faster. However, if tactile motion is initially encoded in a body-centered frame, how does skin-centered perception emerge? One possibility is that the brain applies inverse coordinate transformations, using the available proprioceptive information to remap the body-centered tactile signals into skin-based coordinates. An alternative account is that skin- and body-centered representations are computed in parallel by partially segregated circuits, with task demands determining which representation is preferentially read out. Discriminating between these possibilities will require neural measurements with high temporal and spatial resolution (e.g., local field potentials or single-unit recordings).

In sum, we show that perception of tactile motion is governed by a flexible, task-dependent integration of tactile and proprioceptive signals, with posture information selectively only modulating motion judgments in a Sternum-centric reference frame. In contrast, Finger-centric motion judgments remain largely invariant to changes in arm posture. RT analyses reveal that Sternum-centric judgments are consistently faster, and modeling shows that this advantage reflects lower decision thresholds rather than enhanced accumulation of sensory evidence. Together, these findings indicate that reference frame task demands not only determine how tactile motion signals are transformed by proprioception, but also how those signals are prioritized for perceptual decision making. Further, by introducing a generative framework that moves beyond skin-centered computational models, our work formalizes how proprioceptive signals reshape tactile motion representations to support flexible, reference frame-dependent perception of touch information.

## Acknowledgements

We would like to acknowledge Ms. Amy Fuller and Mr. Yihan Xie for their contribution in collecting data. We also would like to thank Dan Guarino and Martin Gira for their help in building the automated devices. This work was supported with funds from the National Institute of Neurological Disorders and Stroke (NS114191 and NS133935; MGR).

## Author Contributions

**Himanshu Ahuja:** Conceptualization, Methodology, Investigation, Formal analysis, Data Curation and Writing. **Sabyasachi Shivkumar:** Formal analysis and Writing. **Catalina Feistritzer:** Investigation and Data Curation. **Ralf M. Haefner:** Data Curation and Writing. **Gregory C. DeAngelis:** Conceptualization, Methodology, Data Curation and Writing. **Manuel Gomez-Ramirez:** Conceptualization, Methodology, Investigation, Formal analysis, Data Curation, and Writing.

## Data and code availability

All data reported in this paper are stored in Neurodata Without Borders (NWB) format, and will be shared by the corresponding author upon request. Any additional information required to reanalyze the data reported in this paper is available from the corresponding author upon request. It is expected that publications produced from a shared data agreement would result in co-authorships for all of the authors in this manuscript.

