## Supplementary Text and Figures for "Flexible Perception of Tactile Cues in Multiple Reference Frames"

### Supplementary Material and Figures

#### Supplementary Figure 1

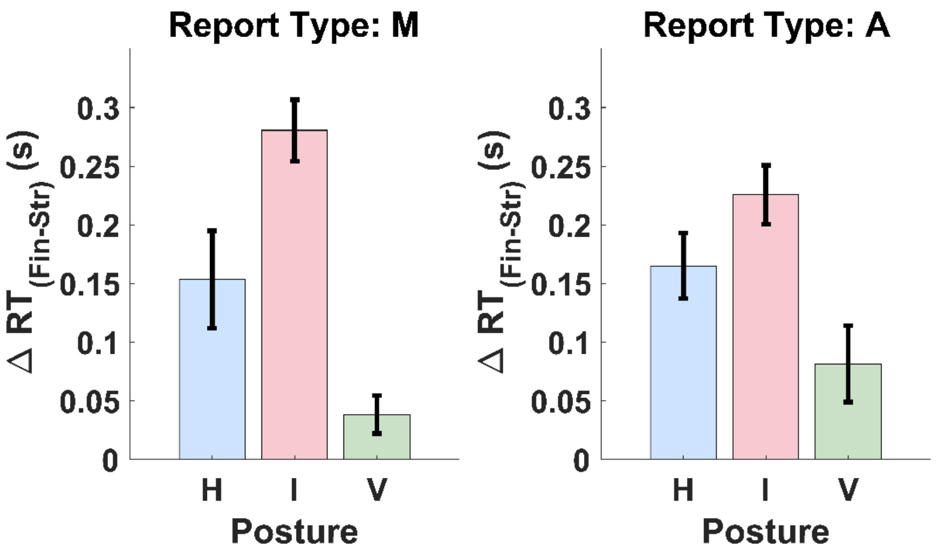


**Supplementary Figure 1**: Difference in Response times (RTs) between the Finger- and Sternum-centric task for the three proprioceptive conditions in Experiment 1. The left and right charts show RTs data in participants responding with a mouse button press (N = 5) and verbal response (N = 7), respectively. The data show very similar pattern of effects across the two response type conditions.

**Supplementary Figure 2**

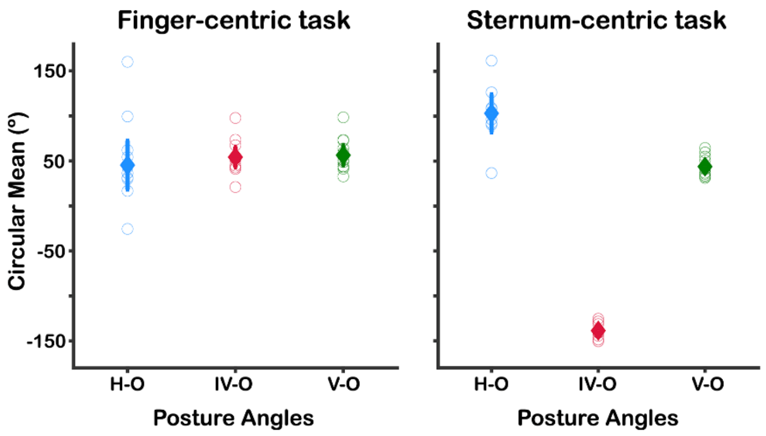


**Supplementary Figure 2:** Circular mean values of each participant for each posture angle in the Finger-centric (left graph) and Sternum-centric (right graph) reference frame task. For each participant, we transformed the psychometric data in each posture and task condition into a vector distribution. The length of each vector equals the proportion of choices in the psychometric curve. The circular mean of each vector distribution was computed to statistically test for posture-dependent shifts in participants' response distributions. Open circles represent the circular mean for each participant. Diamonds represent the circular mean averaged across participants. Error bars denote 95% confidence intervals. N = 14.

This analysis was performed because our experimental paradigm yielded two points of subjective equality (PSE; defined as the angles generating 50% responses) for every psychometric curve. Taking the average across PSE values can yield incorrect values for certain conditions. For example, the circular mean of the two PSE angles in the IV-O posture is ~45º for both Finger- and Sternum-centric conditions. However, as **Figure 2A** (*right panel)* in the main text shows, these two distributions are ~180º out of phase with each other. Transforming the proportion values into vector distributions enable us to compute circular means, and perform circular statistical tests (e.g., Watson-Williams multi sample tests) to determine whether participants' response distributions change as a function of posture.

**Supplementary Figure 3**

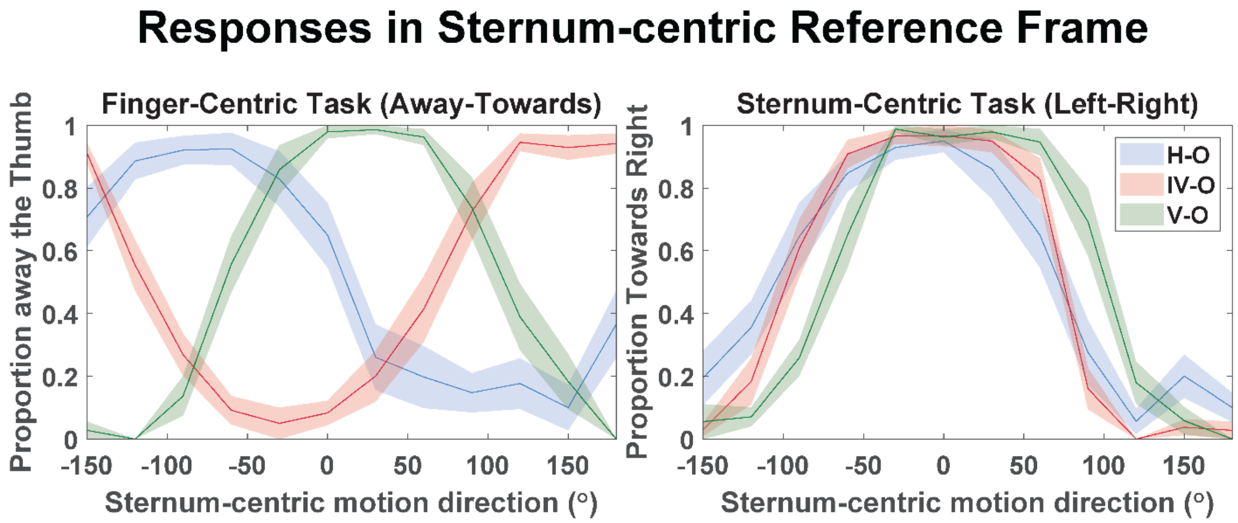


**Supplementary Figure 3:** Population response (N=12) in the Finger-centric task (*left*) and Sternum-centric task (*right*) plotted as a function of stimuli moving in world-centered coordinates. V-O, IV-O, H-O indicate the proprioceptive conditions. Shaded regions represent the standard error of the mean across participants.

**Supplementary Figure 4**

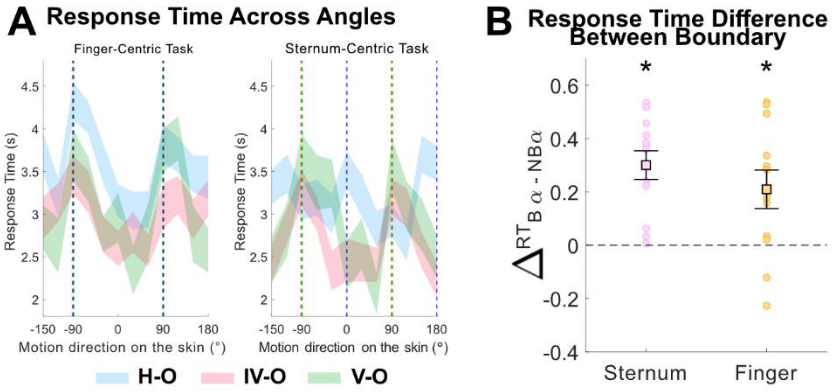


**Supplementary Figure 4A** shows RTs during Finger-centric and Sternum-centric tasks as a function of stimulus motion direction on the skin. A GLM test with factors of angle type (Boundary angles - *B*α; defined as angles within ±Δ30º of the angle that results in 50% perceptual responses, and Non-Boundary angles - *NB*α; defined as motion angles more than ±Δ30º from the angle that results in 50% perceptual responses) and reference frame task showed a main effect of angle type (*B*α vs. *NB*α; F(1,44) = 37.21, $p = 2.4 \times 10^{-7}$), with RTs for *NB*α significantly faster. We did not observe a significant interaction between reference frame and angle type condition (F(1,44) = 1.18, p = 0.28). **Supplementary Figure 4B** shows RT differences between *B*α and *NB*α for each reference frame task condition. These findings indicate that angles around the *B*α are more difficult to discriminate in each task, suggesting that participants performed the tasks in the instructed reference frame.

**Supplementary Figure 5**

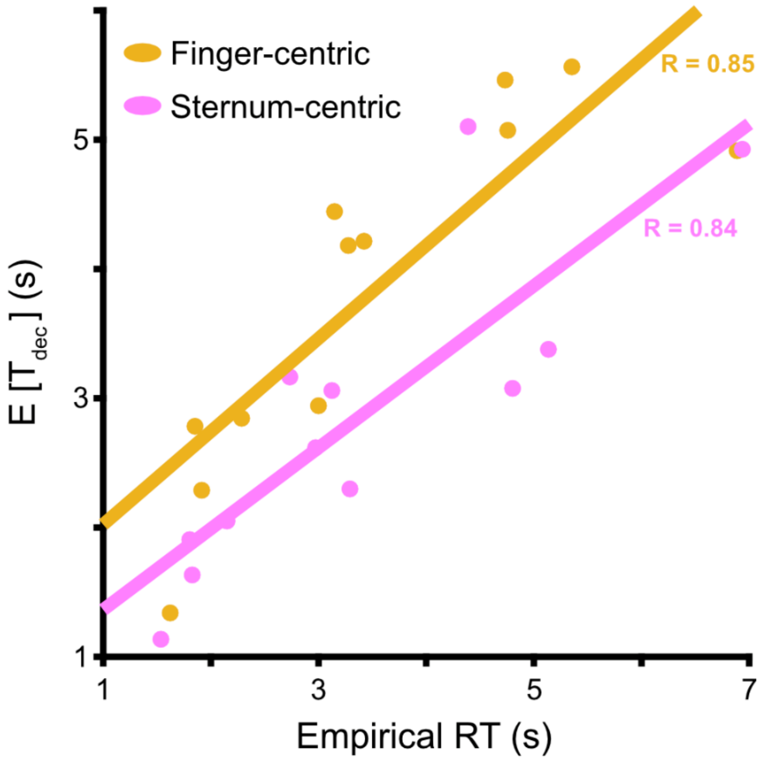


**Supplementary Figure 5**: Expected decision time (E[$T_{dec}$]) of each participant (individual circles) for each reference frame task plotted as a function of the empirical RT for the corresponding reference frame task condition (Finger-centric task = orange data, and Sternum-centric task = magenta data). Expected decision time was computed using Equation 5 in the main text. Expected decision time estimates were strongly correlated with empirical mean RTs across participants in both the Finger-centric task (Pearson correlation, R = 0.85) and the Sternum-centric task (R = 0.84). These results illustrate the nonlinear interaction between drift and decision threshold in shaping response latency, indicating that reduced decision thresholds can offset lower drift gain to produce faster RTs. N = 12. The lines represent linear fits between the empirical and model RT data. The correlation analyses revealed significant relationships across the two RT data for each task (Finger task $p = 4.43 \times 10^{-4}$; Sternum task $p = 5.81 \times 10^{-4}$).

**Supplementary Figure 6**

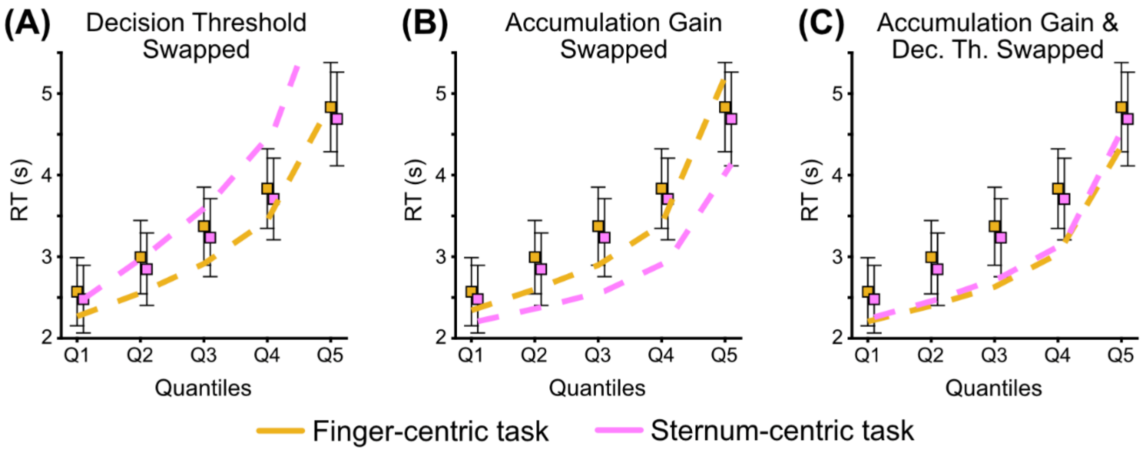


**Supplementary Figure 6:** Plots show mean predicted RTs (model simulations; dashed lines) and empirical mean RTs (boxes) for the Finger-centric and Sternum-centric tasks under different parameter-swapping conditions. We systematically swapped fitted drift gain and/or decision threshold parameters across tasks and generated forward RT predictions from the modified models to determine which parameter drives the faster RTs observed in the Sternum-centric task. **(A)** Swapping decision threshold parameters between the Finger- and Sternum-centric tasks reversed the empirical RT pattern, eliminating the Sternum-centric RT advantage. **(B)** Swapping drift gain parameters produced RT predictions that deviated substantially from the empirical data and failed to reproduce the observed task difference. **(C)** Swapping both drift gain and boundary separation yielded a combination of these effects. Importantly, none of the parameter-swapping combinations reproduced the empirical/DDM RT relationship, which only the non-swapped parameters revealed (see **Figure 2F** in main text). Collectively, these data indicate that the Sternum-centric RT advantage is primarily driven by reduced decision threshold rather than differences in drift gain.

**Supplementary Figure 7**

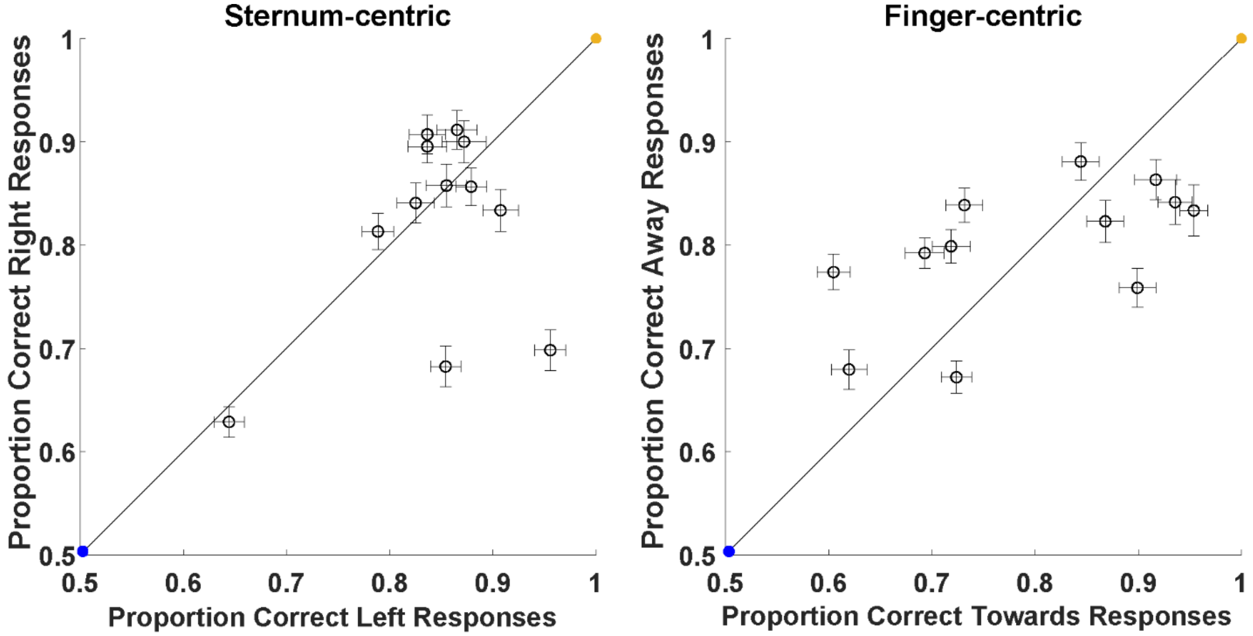


**Supplementary Figure 7**: Choice responses for the Sternum-centric (left plot) and Finger-centric (right plot) reference frame tasks for the behavioral responses in each participant in Experiment 1 (open circles, N = 12). For the Sternum-centric task, the proportion of 'correct right' (or correct left) represents the proportion of trials in which the participant responded 'right' (or 'left') when the stimulus moved rightward (or leftward). Similarly, for the Finger-centric task, the proportion of 'correct away' (or correct 'towards') from the thumb represents the proportion of trials in which the participant responded 'away' ('or 'towards') when the stimulus moved 'away' from (or 'towards') the thumb.

In the Finger-centric task (right plot), correct trials for the 'away' thumb condition were defined as stimuli moving between -90º and +90º. Correct trials for the ''towards' thumb condition were defined as motion stimuli greater than 90º or less than -90º. For the Finger-centric analysis, we used the arrangement of angles shown in the x-axes of **Figure 2A** in the main text. In the Sternum-centric condition (left plot), correct trials for the ‘rightward’ condition were defined as stimuli moving between -90º and +90º. Correct trials for the ‘leftward’ condition were defined as motion stimuli greater than 90º or less than -90º. For the Sternum-centric analysis, we used the arrangement of angles shown in the x-axes of **Supplementary Figure 3**.

Any point on the identity line (black solid line) denotes zero choice preference for Left vs. Right (Sternum-centric task) or Towards vs. Away (Finger-centric task) responses. The orange point (1, 1) denotes no preference for a particular response type, and perfect performance in both tasks. The blue point (0.5, 0.5) denotes no preference for a particular response type, but random behavioral performance in each task. In the Sternum-centric task, a value above vs. below the identity line indicates a preference for choosing ‘Right’ over ‘Left’ responses, respectively. In the Finger-centric task, a value above vs. below the identity line indicate a preference for choosing for choosing ‘Away’ vs. ‘Towards’ the thumb, respectively. The error bars represent standard deviation across 1000 repeated simulations, each with randomly assigned ideal responses to stimulus angles on the discrimination boundary.

### Supplementary Figure 8

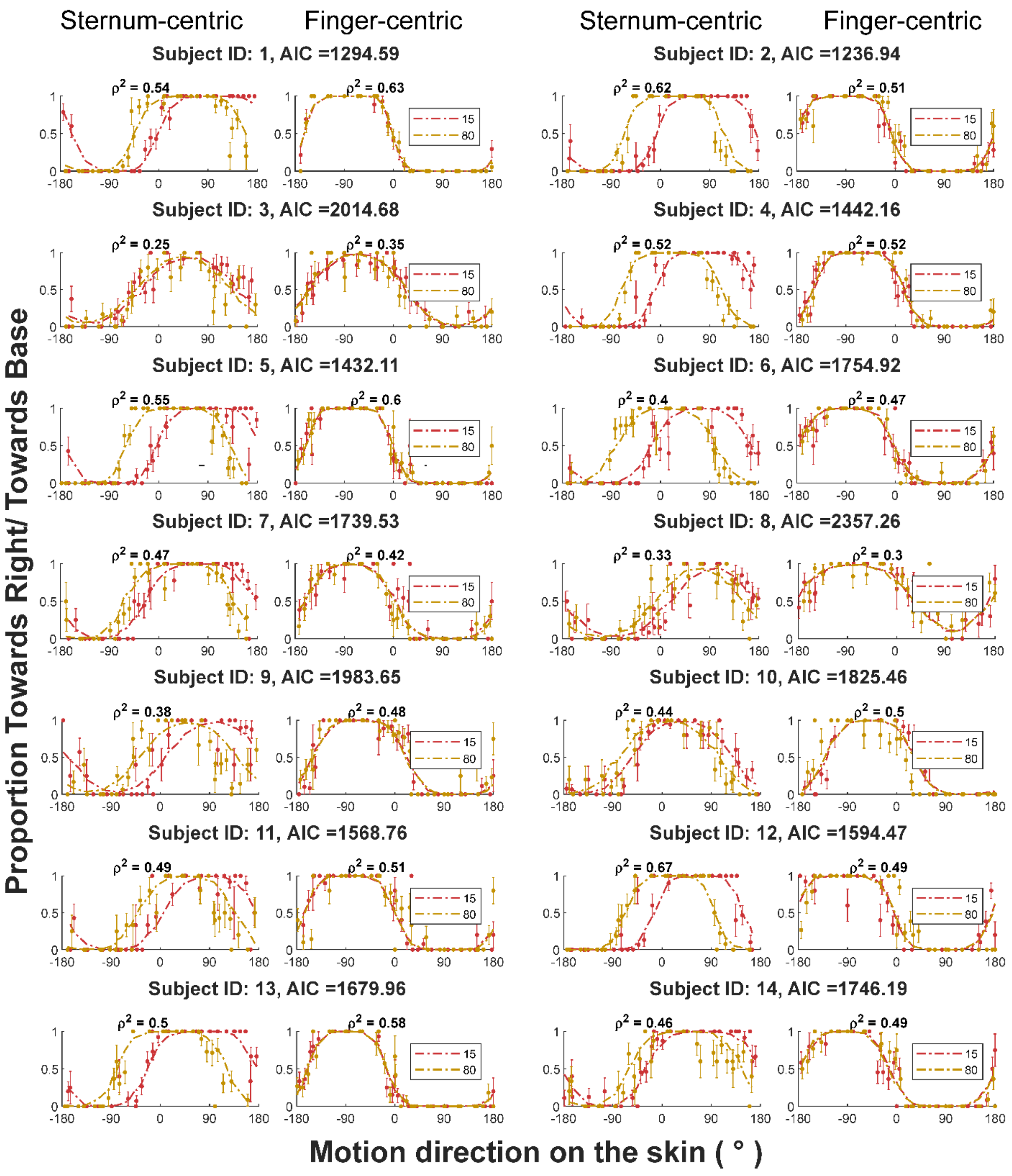

**Supplementary Figure 8**: Psychometric functions for individual participants in Experiment 2. Data are only shown for the two extreme posture angle conditions (15˚ and 80˚, left and right panels, respectively) for visual purposes. Dotted line indicates the fit from the Bayesian Model. McFadden's values are shown for each fit by task, and the AIC score of the model is shown in the title.

**Table 1**

| | McFadden's $\rho^2$ | | Akaike Information Criterion (AIC) | |
|---|---|---|---|---|
| **Participant** | **Euler** | **Bayesian model** | **Euler** | **Bayesian model** |
| 1 | 0.52 | 0.59 | 1504 | 1295 |
| 2 | 0.56 | 0.56 | 1233 | 1237 |
| 3 | 0.28 | 0.30 | 2072 | 2015 |
| 4 | 0.51 | 0.52 | 1469 | 1442 |
| 5 | 0.54 | 0.57 | 1528 | 1432 |
| 6 | 0.44 | 0.44 | 1742 | 1755 |
| 7 | 0.40 | 0.44 | 1872 | 1740 |
| 8 | 0.25 | 0.31 | 2578 | 2357 |
| 9 | 0.43 | 0.43 | 1975 | 1984 |
| 10 | 0.45 | 0.47 | 1911 | 1825 |
| 11 | 0.49 | 0.50 | 1602 | 1569 |
| 12 | 0.57 | 0.58 | 1625 | 1594 |
| 13 | 0.45 | 0.54 | 2014 | 1680 |
| 14 | 0.41 | 0.48 | 1961 | 1746 |
| **Average** | ***0.45*** | ***0.48*** | ***1792*** | ***1691*** |

**Table 1 -** Goodness of fit values for each model fitted to each participant's data in Experiment 2. The $\rho^2$ scores are shown on the left, while the Akaike information criterion (AIC) values are shown on the right. For each participant, the model(s) providing $\Delta$AIC below -20 for the Bayesian model compared to the Euler are highlighted by orange-shaded cells (i.e., significantly better model fits). Relative to the Euler model, the Bayesian model equal or greater $\rho^2$ in all participants, and $\Delta$AIC lower than -20 in 11 out of 14 participants.

### Supplementary Figure 9

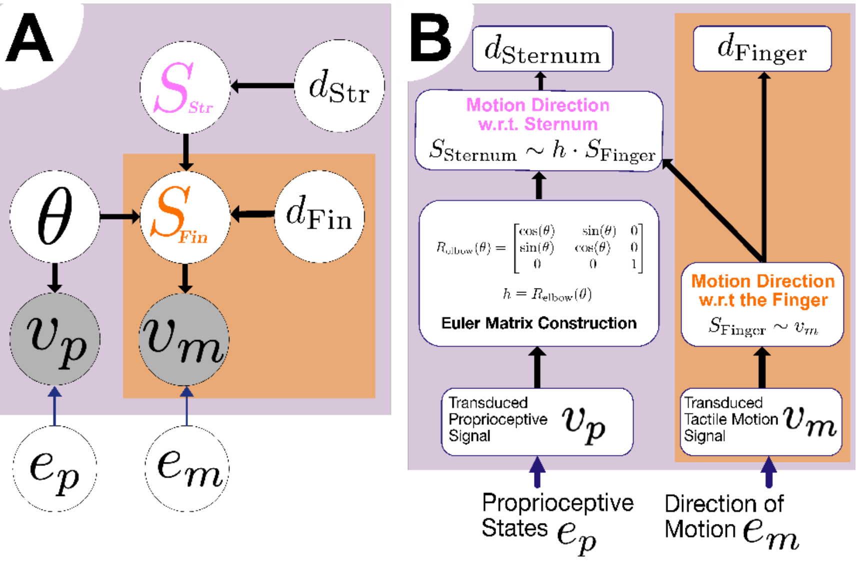


**Supplementary Figure 9:** Conceptual framework of the generative model. Information flow and the generative model for the task. **(A)** A generative model of the psychophysical task is used to make quantitative predictions that can be compared to behavior. The decision variables in the model represent the decisions that the brain makes about the motion direction of the wheel **(Figure 2A inset** in the main text). Here, the decision in a reference frame denotes the models' decision about whether the stimulus is moving left or right in the body-centric reference frame ($d_{\mathrm{Sternum}}$, $d_{\mathrm{str}}$) or towards the tip or base of the stimulated finger in the finger-centric reference frame ($d_{\mathrm{Finger}}$, or $d_{\mathrm{fin}}$). The inferred state variables represent what the brain believes about the current state of environmental parameters, where $S_{\mathrm{Sternum}}$ denotes the perceived direction of motion in the body-centric reference frame, and $S_{\mathrm{Finger}}$ denotes the direction of motion in the Finger-centric reference frame. Finally, the sensory observation $v$ represents the state variable observed through our sensory organs, namely the motion direction on the skin of digit 2 in our task ($v_m$) and the arm posture ($v_p$). $e$ represents the experimental stimulus and $h$ represents the Euler Matrix Transformation from the Finger-centric to the Sternum-centric reference frame. In our task, $h$ is equivalent to rotation at the elbow (**Figure 4A** in main text). **(B)** The information flow has the arrows pointed in the opposite direction to that of the generative model. This distinction stems from the idea that, in the generative model, the sensory observation $v_m$ and $v_p$ is generated from the inferred state variables ($e_m$ and $e_p$).

**Bayesian Generative Modeling**

In this section, we discuss the fundamental generative model underlying the Euler and the Bayesian Generative model with priors on proprioception and perceptual decisions in each task. Each subsequent subsection highlights the modifications on the base model into the respective models mentioned above.

Motion directions in the real world are represented as vectors $v_{x,y,z} \in \mathbb{R}^{3\times1}$. For simplicity of our derivations, we assume a scalar motion direction in the $x - y$ plane represented by $v_m$. The experimenter's generative model produces an observation $v_m$, sampled around the true value $e_m$ with variance $\sigma_m^2$,

$$v_m \sim \mathcal{N}(e_m, \sigma_m^2) \tag{1}$$

and an observation $v_p$, sampled around the true posture $e_p$ with variance $\sigma_p$.

$$v_p \sim \mathcal{N}\left(e_p, \sigma_p^2\right) \tag{2}$$

It's assumed that the brain's (or the observer's) generative model produces the *observed* $v_m$ sampled from a normal distribution with mean $S_{\text{Finger}}$ (*inferred* motion direction in the finger-centric task), and variance $\sigma_m^2$.

$$v_m \sim \mathcal{N}\left(S_{\text{Finger}}, \sigma_m^2\right) \tag{3}$$

The same assumption is held for posture, such that the generative model produces $v_p$ sampled from a normal distribution with mean $\theta$ (inferred posture by the brain in the sternum-centric task), and variance $\sigma_p^2$.

$$v_p \sim \mathcal{N}\left(\theta, \sigma_p^2\right) \tag{4}$$

The generative model has to infer the probability $p\left(d_{\text{Finger}}|e_m\right)$. This can be described by the predictive distribution as follows:

$$p\left(d_{\text{Finger}}|e_m\right) = \int p\left(d_{\text{Finger}}|e_m, v_m\right)p(v_m|e_m)dv_m \tag{5}$$

This marginalization operation can be approximated sampling different values of $v_m$ around $e_m$, and computing $p\left(d_{\text{Finger}}|v_m\right)$.

$$p\left(d_{\text{Finger}}|e_m\right) = \mathbb{E}_{v_m \sim p(v_m|e_m)}\left[p\left(d_{\text{Finger}}|v_m\right)\right] \tag{6}$$

This can be written as,

$$p\left(d_{\text{Finger}}|e_m\right) = \frac{1}{T}\sum_{v_m^{(i)} \sim \mathcal{N}\left(e, \sigma_m^2\right)}^{T}\left[p\left(d_{\text{Finger}}|v_m^{(i)}\right)\right] \tag{7}$$

The predictive distribution for the sternum-centric task is as follows

$$p\left(d_{\text{Sternum}}|e_m, e_p\right) = \int p\left(d_{\text{Sternum}}|e_m, e_p, v_m, v_p\right)p(v_m|e_m)p\left(v_p|e_p\right)dv_m dv_p \tag{8}$$

Similar to the approximation above, now we sample $v_p$ and $v_m$ on each trial. It is important to note that these samples are independent of each other.

$$p\left(d_{\text{Sternum}}|e_m, e_p\right) = \frac{1}{T} \sum_{\substack{v_m^{(i)} \sim \mathcal{N}(e_m, \sigma_m^2) \\ v_p^{(i)} \sim \mathcal{N}(e_p, \sigma_p^2)}}^{T} \left[p\left(d_{\text{Sternum}}|v_m^{(i)}, v_p^{(i)}\right)\right] \tag{9}$$

For brevity of notation, we implicitly assume the subscript over $v_m$ and $v_p$ across the $T$ trials. A decision for an experimental condition (for instance, in the finger-centric condition) is made by comparing the posterior probabilities $p(d_{\text{Finger}} = t|v_m)$ and $p(d_{\text{Finger}} = a|v_m)$.

$$d_{\text{Finger}} = \begin{cases} 1, & \text{if } p\left(d_{\text{Finger}} = a|v_m\right) > p\left(d_{\text{Finger}} = t|v_m\right) \\ 0, & \text{otherwise} \end{cases} \tag{10}$$

Here, the binary variable $d_{\text{Finger}}$ takes the value 1 if the probability that the stimulus is moving away from the thumb $(a)$ is greater than the probability of stimulus moving towards the thumb $(t)$.

Across multiple trials $T$, we can then calculate the probability $p(d_{\text{Finger}} = a|e_m)$ as:

$$p\left(d_{\text{Finger}} = a|e_m\right) = \frac{1}{T} \sum_{v_m \sim \mathcal{N}(e_m, \sigma_m^2)}^{T} \mathbb{I}\left(\frac{p\left(d_{\text{Finger}} = a|v_m\right)}{p\left(d_{\text{Finger}} = t|v_m\right)} > 1\right) \tag{11}$$

The posterior probability $p(d_{\text{Finger}} = a|v_m)$ is expanded using Bayes' rule

$$p\left(d_{\text{Finger}} = a|v_m\right) = \frac{p\left(v_m|d_{\text{Finger}} = a\right)p\left(d_{\text{Finger}} = a\right)}{p(v_m)} \tag{12}$$

Participants might be biased towards making one choice over the other. We parameterize this choice bias in the finger-centric reference frame as,

$$p\left(d_{\text{Finger}} = a\right) = \Gamma_{\text{Finger}} \tag{13}$$

and in the sternum-centric reference frame as follows:

$$p(d_{\text{Sternum}} = r) = \Gamma_{\text{Sternum}} \tag{14}$$

We can now re-write Equation 11 substituting from Equation 12 and Equation 13.

$$p(d_{\text{Finger}} = a | e_m) = \frac{1}{T} \sum_{v_m \sim \mathcal{N}(e_m, \sigma_m^2)}^{T} \mathbb{I}\left( \frac{p(v_m | d_{\text{Finger}} = a) \cdot \Gamma_{\text{Finger}}}{p(v_m | d_{\text{Finger}} = t) \cdot (1 - \Gamma_{\text{Finger}})} > 1 \right) \tag{15}$$

Similarly, for the sternum-centric task, we can write

$$p(d_{\text{Sternum}} = r | e_m, e_p) = \frac{1}{T} \sum_{\substack{v_m \sim \mathcal{N}(e_m, \sigma_m^2) \\ v_p \sim \mathcal{N}(e_p, \sigma_p^2)}}^{T} \mathbb{I}\left( \frac{p(v_m, v_p | d_{\text{Sternum}} = r) \cdot \Gamma_{\text{Sternum}}}{p(v_m, v_p | d_{\text{Sternum}} = l) \cdot (1 - \Gamma_{\text{Sternum}})} > 1 \right) \tag{16}$$

The likelihood term, $p(v_m | d_{\text{Finger}} = a)$ can be written as a marginalization over the intermediate variable $S_{\text{Finger}}$.

$$p(v_m | d_{\text{Finger}} = a) = \int p(v_m | S_{\text{Finger}}) p(S_{\text{Finger}} | d_{\text{Finger}} = a) dS_{\text{Finger}} \tag{17}$$

where $p(v | S_{\text{Finger}})$ is represented using a normal distribution with variance parameter $\sigma_m^2$,

$$p(v_m | S_{\text{Finger}}) = \frac{1}{\sigma_m \sqrt{2\pi}} e^{-\frac{1}{2}(\frac{v_m - S_{\text{Finger}}}{\sigma_m})^2} \tag{18}$$

and the conditional $p(S_{\text{Finger}} | d_{\text{Finger}} = a)$ as a circular step-function, at the decision boundary at $\ldots, -\pi, \pi, 3\pi, \ldots$ radians.

$$p(S_{\text{Finger}} | d_{\text{Finger}} = a) = H(S_{\text{Finger}} - (2k + 2)\pi) - H(S_{\text{Finger}} - (2k + 1)\pi), k \in \mathbb{Z} \tag{19}$$

Populating Equations 19 and 18 in Equation 17:

$$p\left(v_m|d_{\text{Finger}}=a\right)=\int \mathcal{N}\left(v_m;S_{\text{Finger}},\sigma_m^2\right)\left[H\left(S_{\text{Finger}}\right)-H\left(S_{\text{Finger}}-\pi\right)\right]\mathrm{d}S_{\text{Finger}} \tag{20}$$

$$p\left(v_m|d_{\text{Finger}}=a\right)=\Phi(\pi;v_m,\sigma_m^2)-\Phi(0;v_m,\sigma_m^2) \tag{21}$$

In the above equation, $\Phi$ represents the cumulative Gaussian function. Hence, for the finger-centric reference frame the response function is:

$$d_{\text{Finger}}=\begin{cases}1, & \text{if} \quad \dfrac{\Phi(\pi;v_m,\sigma_m^2)-\Phi(0;v_m,\sigma_m^2)}{\Phi(2\pi;v_m,\sigma_m^2)-\Phi(\pi;v_m,\sigma_m^2)}>\dfrac{\left(1-\Gamma_{\text{Finger}}\right)}{\Gamma_{\text{Finger}}}\\ 0, & \text{otherwise}\end{cases} \tag{22}$$

Similar to Equation 17, $p\left(v_m,v_p|d_{\text{Sternum}}=r\right)$ can be written as a marginalization over the intermediate variables $S_{\text{Sternum}}$ and $S_{\text{Finger}}$.

$$p\left(v_m,v_p|d_{\text{Sternum}}=r\right)=\int\int\int p\left(v_m|S_{\text{Finger}}\right)p\left(S_{\text{Finger}}|S_{\text{Sternum}}\right)p\left(v_p|\theta\right) p(S_{\text{Sternum}}|d_{\text{Sternum}}=r)\mathrm{d}S_{\text{Finger}}\mathrm{d}S_{\text{Sternum}}\mathrm{d}\theta \tag{23}$$

Note the additional term $p\left(S_{\text{Finger}}|S_{\text{Sternum}}\right)$ in the equation. This term represents the probability distribution of the transformation from the finger-centric reference frame to the sternum-centric reference frame. We assume this distribution to be normal with mean given by $q^{-1}\left(h\cdot q\left(S_{\text{Finger}}\right)\right)$ and variance $\sigma_h^2$.

$$p\left(S_{\text{Finger}}|S_{\text{Sternum}}\right)=\frac{1}{\sigma_h\sqrt{2\pi}}e^{-\frac{1}{2}\left(\frac{q^{-1}\left(h\cdot q\left(S_{\text{Finger}}\right)\right)-S_{\text{Sternum}}}{\sigma_h}\right)^2} \tag{24}$$

Here, $h$ is the Euler Matrix transformation from the finger-centric reference frame to the sternum-centric reference frame.

$$h=R_{\text{elbow},z}(\theta) \tag{25}$$

Here, $R_{\text{elbow},z}$ is the rotation by $\theta$ at the elbow, around the $z$-axis.

$$R_{\text{elbow},z}(\theta) = \begin{bmatrix} \cos(\theta) & -\sin(\theta) & 0 \\ \sin(\theta) & \cos(\theta) & 0 \\ 0 & 0 & 1 \end{bmatrix} \tag{26}$$

and the function $q: \mathbb{R} \rightarrow \mathbb{R}^{3\times1}$ vectorizes the scalar motion direction.

$$q(\gamma) = \begin{bmatrix} \cos(\gamma) \\ \sin(\gamma) \\ 0 \end{bmatrix}, \gamma \in [0,2\pi) \tag{27}$$

In the current experiment, we only work with linear transformations, which implies that the Euler Matrix transformation can simply be represented as an additive term, i.e.

$$q^{-1}\left(R_{\text{elbow},z}(\theta) \cdot q\left(S_{\text{Finger}}\right)\right) \equiv S_{\text{Finger}} + \theta \tag{28}$$

The transformation can further be generalized to other reference frames as a product of rotation matrices.

$$h = R_{\text{elbow},z}(\theta_1) \cdot R_{\text{shoulder},z}(\theta_2) \cdot R_{\text{elbow},x}(\theta_3) \cdot \ldots \tag{29}$$

Similar to the step function described above (Equation 19), the discrimination boundary (in this task, it is at $\pi$), for the sternum-centric task is described as:

$$p(S_{\text{Sternum}}|d_{\text{Sternum}} = r) = H\left(S_{\text{Sternum}} - (2k)(\pi)\right)) - H\left(S_{\text{Sternum}} - (2k+1)(\pi)\right), k \in \mathbb{Z} \tag{30}$$

Populating Equations 30, 28 and 24 in Equation 23:

$$p\left(v_m, v_p|d_{\text{Sternum}} = r\right) = \int\int\int \mathcal{N}\left(v_m; S_{\text{Finger}}, \sigma_m^2\right)\mathcal{N}\left(S_{\text{Finger}}; S_{\text{Sternum}} - \theta, \sigma_h^2\right) \mathcal{N}\left(v_p; \theta, \sigma_p^2\right)[H(S_{\text{Sternum}}) - H(S_{\text{Sternum}} - \pi)]\mathrm{d}S_{\text{Finger}}\mathrm{d}S_{\text{Sternum}}\mathrm{d}\theta \tag{31}$$

$$p\left(v_m, v_p | d_{\text{Sternum}} = r\right) = \int\int \mathcal{N}(S_{\text{Sternum}}; v_m + \theta, \sigma_m^2 + \sigma_h^2)\mathcal{N}\left(v_p; \theta, \sigma_p^2\right) \\ [H(S_{\text{Sternum}}) - H(S_{\text{Sternum}} - \pi)]\mathrm{d}S_{\text{Sternum}}\mathrm{d}\theta \tag{32}$$

$$p\left(v_m, v_p | d_{\text{Sternum}} = r\right) = \int \mathcal{N}\left(S_{\text{Sternum}}; v_m + v_p, \sigma_m^2 + \sigma_h^2 + \sigma_p^2\right) \\ [H(S_{\text{Sternum}}) - H(S_{\text{Sternum}} - \pi)]\mathrm{d}S_{\text{Sternum}} \tag{33}$$

$$p\left(v_m, v_p | d_{\text{Sternum}} = r\right) = [\Phi\left(\pi; v_m + v_p, \sigma_m^2 + \sigma_h^2 + \sigma_p^2\right) - \\ \Phi\left(0; v_m + v_p, \sigma_m^2 + \sigma_h^2 + \sigma_p^2\right)] \tag{34}$$

Finally, the response function for the sternum-centric task is:

$$d_{\text{Sternum}} = \begin{cases} 1, & \text{if} \quad \dfrac{\begin{bmatrix} \Phi\left(\pi; v_m + v_p, \sigma_m^2 + \sigma_h^2 + \sigma_p^2\right) - \\ \Phi\left(0; v_m + v_p, \sigma_m^2 + \sigma_h^2 + \sigma_p^2\right) \end{bmatrix}}{\begin{bmatrix} \Phi\left(2\pi; v_m + v_p, \sigma_m^2 + \sigma_h^2 + \sigma_p^2\right) - \\ \Phi\left(\pi; v_m + v_p, \sigma_m^2 + \sigma_h^2 + \sigma_p^2\right) \end{bmatrix}} > \dfrac{(1 - \Gamma_{\text{Sternum}})}{\Gamma_{\text{Sternum}}} \\ 0, & \text{otherwise} \end{cases} \tag{35}$$

**Euler model**

We consider the Euler-matrix transformation to be exact, such that $\sigma_h^2 \to 0$. This was done because our current experimental paradigm cannot separate transformation noise from proprioceptive noise. So, we use $\sigma_p^2$ as a placeholder for the total variance under transformation and proprioception. Secondly, we assumed that the subjects have no prior bias in making a decision towards or away from the thumb in the finger-centric reference frame or in making a decision left or right the center of the body in the sternum-centric reference frame. This implies that $\Gamma_{\text{Finger}} = 0.5$, and $\Gamma_{\text{Sternum}} = 0.5$. Lastly, we add a bias term on the perceived motion direction in each reference frame: $\beta_{\text{Finger}}$ for the finger-centric reference frame and $\beta_{\text{Sternum}}$ for the sternum-centric reference frame.

With the above modifications, the response function for the finger-centric task becomes,

$$d_{\text{Finger}} = \begin{cases} 1, & \text{if} \quad \dfrac{\Phi\left(\pi; v_m + \beta_{\text{Finger}}, \sigma_m^2\right) - \Phi\left(0; v_m + \beta_{\text{Finger}}, \sigma_m^2\right)}{\Phi\left(2\pi; v_m + \beta_{\text{Finger}}, \sigma_m^2\right) - \Phi\left(\pi; v_m + \beta_{\text{Finger}}, \sigma_m^2\right)} > 1 \\ 0, & \text{otherwise} \end{cases} \tag{36}$$

and for the sternum-centric reference frame,

$$d_{\text{Sternum}} = \begin{cases} 1, & \text{if} \quad \dfrac{\begin{bmatrix} \Phi\left(\pi; v_m + v_p + \beta_{\text{Sternum}}, \sigma_m^2 + \sigma_p^2\right) - \\ \Phi\left(0; v_m + v_p + \beta_{\text{Sternum}}, \sigma_m^2 + \sigma_p^2\right) \end{bmatrix}}{\begin{bmatrix} \Phi\left(2\pi; v_m + v_p + \beta_{\text{Sternum}}, \sigma_m^2 + \sigma_p^2\right) - \\ \Phi\left(\pi; v_m + v_p + \beta_{\text{Sternum}}, \sigma_m^2 + \sigma_p^2\right) \end{bmatrix}} > 1 \\ 0, & \text{otherwise} \end{cases} \tag{37}$$

In total, this model has four parameters: two variance parameters, $\sigma_m^2$ and $\sigma_p^2$, and two bias parameters, $\beta_{\text{Finger}}$ and $\beta_{\text{Sternum}}$. With each new reference frame condition, this model will simply require two additional parameters. The parameters of this model do not scale with the number of joint transformations, or rotational axes.

**Bayesian Generative Model with Prior on Proprioception and Choice**

The finger-centric response function in this condition includes a decision bias parameter compared to the ideal observer model, such that Equation 22 becomes,

$$d_{\text{Finger}} = \begin{cases} 1, & \text{if} \quad \dfrac{\Phi\left(\pi; v_m + \beta_{\text{Finger}}, \sigma_m^2\right) - \Phi\left(0; v_m + \beta_{\text{Finger}}, \sigma_m^2\right)}{\Phi\left(2\pi; v_m + \beta_{\text{Finger}}, \sigma_m^2\right) - \Phi\left(\pi; v_m + \beta_{\text{Finger}}, \sigma_m^2\right)} > \dfrac{\left(1 - \Gamma_{\text{Finger}}\right)}{\Gamma_{\text{Finger}}} \\ 0, & \text{otherwise} \end{cases} \tag{38}$$

For the sternum-centric condition, we assume a prior distribution over posture $\theta$ $p\left(\theta | \mu, \sigma_\mu^2\right) = \mathcal{N}\left(\theta; \mu, \sigma_\mu^2\right)$. This modifies Equation 23, such that the likelihood is given by,

$$p\left(v_m, v_p | d_{\text{Sternum}} = r\right) = \int\int\int p\left(v_m | S_{\text{Finger}}\right) p\left(S_{\text{Finger}} | S_{\text{Sternum}}, \theta\right) p\left(v_p | \theta\right) \\ p(\theta|\mu) p(S_{\text{Sternum}} | d_{\text{Sternum}} = r) \mathrm{d}S_{\text{Sternum}} \mathrm{d}S_{\text{Finger}} \mathrm{d}\theta \tag{39}$$

Next, we apply a series of substitutions, similar to the ones performed in Equations 24 to 34:

$$p\left(v_m, v_p | d_{\text{Sternum}} = r\right) = \int\int\int \mathcal{N}\left(v_m; S_{\text{Finger}}, \sigma_m^2\right) \mathcal{N}\left(S_{\text{Finger}}; S_{\text{Sternum}} - \theta; \sigma_h^2\right) \\ \mathcal{N}\left(v_p; \theta, \sigma_p^2\right) \mathcal{N}\left(\theta; \mu, \sigma_\mu^2\right) \\ [H(S_{\text{Sternum}}) - H(S_{\text{Sternum}} - \pi)] \mathrm{d}S_{\text{Sternum}} \mathrm{d}S_{\text{Finger}} \mathrm{d}\theta \tag{40}$$

$$p\left(v_m, v_p | d_{\text{Sternum}} = r\right) = \int\int \mathcal{N}(v_m; S_{\text{Sternum}} - \theta, \sigma_h^2 + \sigma_m^2) \\ \mathcal{N}\left(v_p; \mu, \sigma_p^2 + \sigma_\mu^2\right) \mathcal{N}\left(\theta; \frac{\mu\sigma_p^2 + v_p\sigma_\mu^2}{\sigma_p^2 + \sigma_\mu^2}, \frac{\sigma_p^2\sigma_\mu^2}{\sigma_p^2 + \sigma_\mu^2}\right) \\ [H(S_{\text{Sternum}}) - H(S_{\text{Sternum}} - \pi)] \mathrm{d}S_{\text{Sternum}} \mathrm{d}\theta \tag{41}$$

$$p\left(v_m, v_p | d_{\text{Sternum}} = r\right) = \int \mathcal{N}\left(S_{\text{Sternum}}; v_m + \frac{\mu\sigma_p^2 + v_p\sigma_\mu^2}{\sigma_p^2 + \sigma_\mu^2}, \sigma_h^2 + \sigma_m^2 + \frac{\sigma_p^2\sigma_\mu^2}{\sigma_p^2 + \sigma_\mu^2}\right) \\ \mathcal{N}\left(v_p; \mu, \sigma_p^2 + \sigma_\mu^2\right) \\ [H(S_{\text{Sternum}}) - H(S_{\text{Sternum}} - \pi)] \mathrm{d}S_{\text{Sternum}} \tag{42}$$

$$p\left(v_m, v_p | d_{\text{Sternum}} = r\right) = [\Phi\left(\pi; v_m + \frac{\mu\sigma_p^2 + v_p\sigma_\mu^2}{\sigma_p^2 + \sigma_\mu^2}, \sigma_h^2 + \sigma_m^2 + \frac{\sigma_p^2\sigma_\mu^2}{\sigma_p^2 + \sigma_\mu^2}\right) - \\ \Phi\left(0; v_m + \frac{\mu\sigma_p^2 + v_p\sigma_\mu^2}{\sigma_p^2 + \sigma_\mu^2}, \sigma_h^2 + \sigma_m^2 + \frac{\sigma_p^2\sigma_\mu^2}{\sigma_p^2 + \sigma_\mu^2}\right)] \\ \mathcal{N}\left(v_p; \mu, \sigma_p^2 + \sigma_\mu^2\right) \tag{43}$$

$$d_{\text{Sternum}} = \begin{cases} 1, & \text{if } \dfrac{\left[\begin{array}{c} \Phi\left(\pi; v_m + \dfrac{\mu\sigma_p^2 + v_p\sigma_\mu^2}{\sigma_p^2 + \sigma_\mu^2}, \sigma_h^2 + \sigma_m^2 + \dfrac{\sigma_p^2\sigma_\mu^2}{\sigma_p^2 + \sigma_\mu^2}\right) - \\ \Phi\left(0; v_m + \dfrac{\mu\sigma_p^2 + v_p\sigma_\mu^2}{\sigma_p^2 + \sigma_\mu^2}, \sigma_h^2 + \sigma_m^2 + \dfrac{\sigma_p^2\sigma_\mu^2}{\sigma_p^2 + \sigma_\mu^2}\right) \end{array}\right]}{\left[\begin{array}{c} \Phi\left(2\pi; v_m + \dfrac{\mu\sigma_p^2 + v_p\sigma_\mu^2}{\sigma_p^2 + \sigma_\mu^2}, \sigma_h^2 + \sigma_m^2 + \dfrac{\sigma_p^2\sigma_\mu^2}{\sigma_p^2 + \sigma_\mu^2}\right) - \\ \Phi\left(\pi; v_m + \dfrac{\mu\sigma_p^2 + v_p\sigma_\mu^2}{\sigma_p^2 + \sigma_\mu^2}, \sigma_h^2 + \sigma_m^2 + \dfrac{\sigma_p^2\sigma_\mu^2}{\sigma_p^2 + \sigma_\mu^2}\right) \end{array}\right]} > \dfrac{(1 - \Gamma_{\text{Sternum}})}{\Gamma_{\text{Sternum}}} \\ 0, & \text{otherwise} \end{cases} \quad (44)$$

In total, this model has 8 parameters. Three parameters for the finger-centric reference frame: $\Gamma_{\text{Finger}}$, $\beta_{\text{Finger}}$ and $\sigma_m^2$, plus five parameters for the sternum-centric reference frame: $\Gamma_{\text{Sternum}}$, $\sigma_p^2$, transformation noise $\sigma_h^2$, mean and variance of the prior over proprioception $\mu$ and $\sigma_\mu^2$. The Bayesian model can explain the posture-dependent changes in circular mean responses of the Sternum-centric task (i.e., the deviation of the slope from -1 that is observed in **Figure 4C** in main text) by a non-uniform prior over arm position. The slope predicted by the model is given by Equation 45:

$$Model\ Predicted\ Slope = -\frac{\sigma_\mu^2}{\sigma_p^2 + \sigma_\mu^2} \quad (45)$$

**Supplementary Figure 10** shows that the slope predicted by the model is in good agreement with participants' empirical slope values.

### Supplementary Figure 10

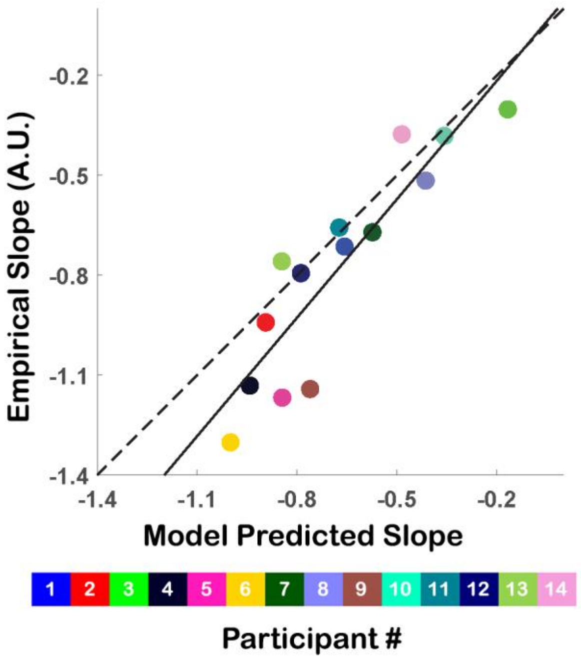


**Supplementary Figure 10:** Comparisons between the Bayesian model's predicted slope and the observed slope of the circular mean in the Sternum-centric task. The match is excellent for empirical slopes between zero and -1. The model cannot in principle explain slopes less than -1 since our unimodal prior over arm position always leads to an under-correction, but never an overcorrection for arm position in the Sternum-centric task.

### Model Fitting

The likelihood of the model parameters $P$ across all experimental conditions - reference frames $RF$, postures $e_p$ and motion directions $e_m$, for $T$ trials per condition is given by the binomial probability,

$$L(P|\text{Data}) = \sum_{i}^{\text{RF}} \sum_{j}^{e_p} \sum_{k}^{e_m} p\left(d_{\text{RF}}^{(i)} | e_m^{(k)}, e_p^{(j)}\right)^t \left(1 - p\left(d_{\text{RF}}^{(i)} | e_m^{(k)}, e_p^{(j)}\right)\right)^{T-t} \quad (46)$$

Here $t$ is the number of responses for choice one, and $T - t$ are the number of responses for choice two. We minimize the negative log likelihood of the parameters to get the best fit of the model using the BADS algorithm[67] in MATLAB (R2021a, Mathwork Inc.). The optimization procedure was repeated 20 times with different parameter initializations, and the final parameters were largely insensitive to initial values.

### Curve Fitting

Similar to the model fitting process, we use the BADS algorithm to minimize the negative log-likelihood of individual psychometric curves at each posture and reference frame combination. Each curve is defined by two parameters, variance ($\sigma^2$) and bias ($\beta$). This curve fitting was used to calculate the perceptual biases of participants (**Figure 4B** in main text).
